# Unified CEBRA learns shared neural latents across recordings to map single-neuron contributions to population codes

**DOI:** 10.64898/2026.11.18.767506

**Authors:** Célia Benquet, Hossein Mirzaei, Steffen Schneider, Mackenzie Weygandt Mathis

## Abstract

Neural computation is thought to rely on distributed populations; however, most single recording samples capture only a fraction of them, and rare cell types may contribute only a small proportion of neurons per session. Pooling many recordings could reveal how these populations encode information, but each recording samples different neurons, so their contributions cannot be easily compared directly. Here we show that recordings can be combined into a single model of all recorded neurons by aligning them with a shared label, used at both training and inference, placing every neuron’s contribution to the population code on a common scale. This approach, called Unified CEBRA, retained interpretable task information across recordings in the monkey and mouse sensorimotor cortex, rat hippocampus, and mouse visual cortex. Pooling thousands of excitatory and inhibitory neurons from hundreds of sessions from mouse visual cortex, we found that the neurons that contributed most to the model’s readout of visual features were not only the most strongly tuned, but also those that responded more reliably and were less coupled to co-recorded neurons. Unified CEBRA, therefore, can unveil coding principles studied at the population scale with mappable features to individual neurons from sparse, multi-session recordings using our new approach.

## Introduction

The brain computes with large populations (1–5), yet most experiments record tens to hundreds of neurons at a time (6, 7). Large-scale recording remains expensive and restricted to a few laboratories (8, 9), while many smaller recordings are openly available (10–13). Pooled, they could reveal population codes that no single experiment captures, including the contribution of molecularly defined cell classes, such as inhibitory interneurons, that are sparse in any one session.

Existing latent variable models capture population structure (14, 15), but are identifiable only when conditioned on an observed auxiliary variable such as behavior or time (16– 19), and models built on this principle treat each recording separately (20–23). Alignment methods match the low-dimensional geometry of recordings (24–33), typically against a reference session that bounds the alignment, or pair-wise at a combinatorial cost (34), or in a shared space that discards neuron identity (35). Stitching and multi-session transformers learn session-specific input layers or unit embeddings (36–42).

This matters because the question is increasingly which neurons drive the different dimensions in a population code. Predictive “digital twins” compare stimulus tuning across recordings, but their neuron readouts are defined relative to a core that differs between training runs (44– Large decoding transformers, a step towards foundation models of neural activity (47, 48), are costly to train and can exploit shortcuts such as session-specific noise or confounds unrelated to neural activity (49). Interpreting them through attention weights (50) or sparse autoencoders (51) lacks identifiability, since attention weights do not need to reflect a unit’s importance (52), and removing units from a trained network introduces out-of-distribution artifacts (53).

Here we introduce *Unified CEBRA*, which trains one encoder on all input neurons. At each timepoint, every session contributes its activity at the matching value of a shared auxiliary variable, such as behavior, trial time or a stimulus. The matched activities are concatenated into a single pseudo-population and embedded with CEBRA’s contrastive objective, so the identifiability guarantees of CEBRA apply (17, 21, 54, 55). Chaining the encoder’s Jacobian with a linear feature readout, rather than inverting it as in xCE-BRA (56, 57), yields a signed neuron-to-feature attribution map that is invariant to the linear transformation relating across training runs and places every neuron on one scale.

Across motor, navigation and visual datasets, the unified embedding retained task information, scaling to hundreds of recording sessions at a fraction of the computational cost of multi-session transformers. It separated behavioral variables into distinct subspaces and could be recovered when individual animals sampled only part of the behavioral space. For example, when applied to 2,924 excitatory and inhibitory neurons from 234 sessions of the Allen Brain Observatory (13), the attribution map was reproducible across runs, and removing its top-ranked neurons specifically impaired decoding of the corresponding feature. The neurons that mattered most were largely the most strongly tuned, but beyond tuning, the map favored neurons that were more reliable and less coupled to their neighbors.

### Unified encoders via label-guided alignment

Recordings from different sessions and animals typically do not share the same neurons or number of neurons, but they often share a task structure, either temporal or behavioral. We use this shared structure to align timepoints across recordings via strict nearest-neighbor matching. Concretely, we introduce a novel sampling strategy to CEBRA (21) that jointly selects reference, positive, and negative samples across recordings (Methods; Figure 1a). Neural activity is matched across recordings guided by labels, concatenated (Figure 1b) and passed through a single encoder trained by a contrastive InfoNCE loss (21, 58) (Figure 1c). The same label-guided assembly of activity is performed during inference using the target session. The resulting trajectory therefore embeds the pooled population, not the target session alone, along that session’s auxiliary variable timeline.

**Figure 1.**
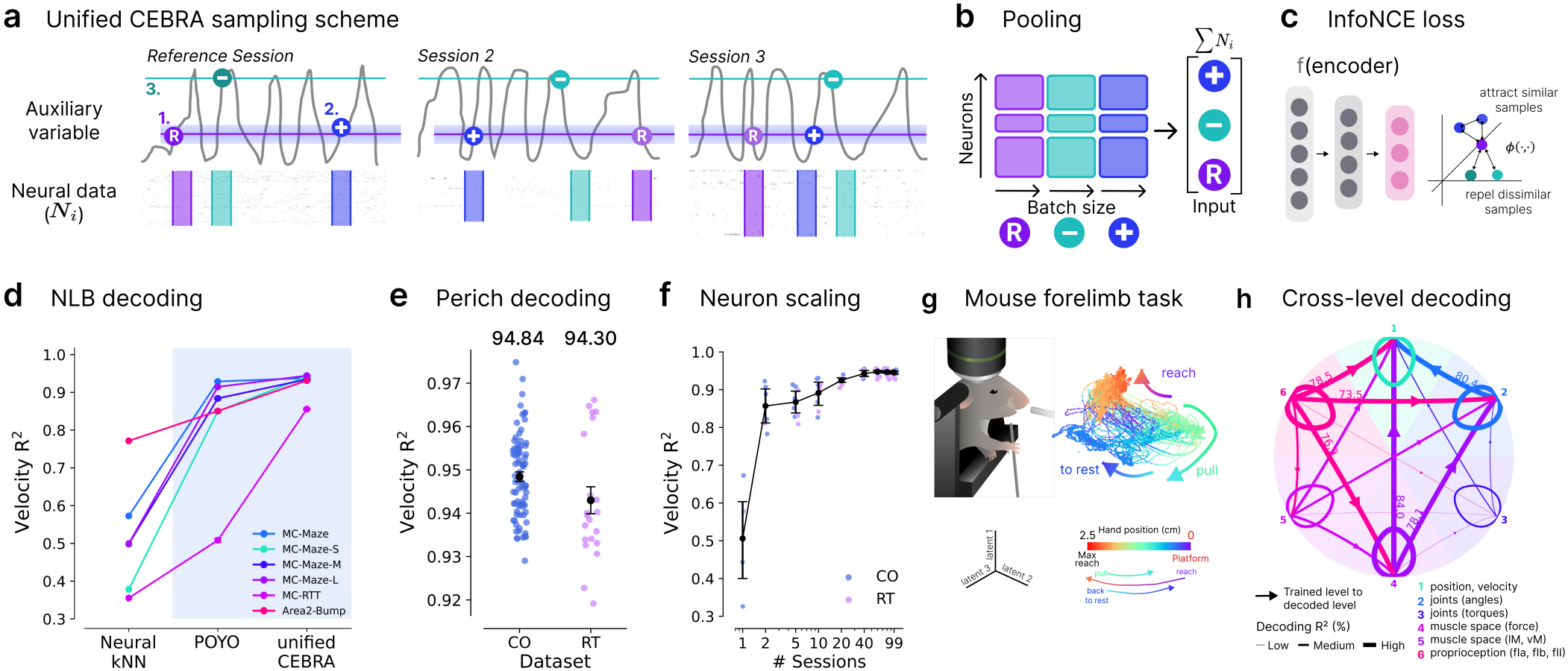
Unified CEBRA pools recordings through a single encoder. **a:** Cross-session sampling with auxiliary labels. *1*.References are drawn uniformly in a reference session and matched in the other sessions by label similarity. *2*. Positives are drawn within each session, close to the reference in label space. *3*. Negatives are drawn like references. **b:** Pooling. At training, references, positives, and negatives are each concatenated across recordings into a “pseudo-session” input; at inference, the target session’s labels select the timepoint of each other session concatenated with the corresponding target timepoint. Labels index the input but are not part of it. **c:** InfoNCE loss for continuous auxiliary variables: positives are attracted to references and negatives are repelled in the latent space. **d:** Velocity decoding across NLB datasets (mean *±*SEM, n = 10 runs for CEBRA variants; public validation data). Blue shading: multi-session methods; values in Supplemental Table S5. **e:** Velocity decoding (R^2^ averaged over x and y; successful reaching held-out data; n = 10 runs) for CO and RT. Colored dots: sessions; black dot: mean*±* SEM. **f:** Velocity decoding as a function of the number of pooled sessions (up to 99 sessions, 9,725 neurons; R^2^, mean *±*SEM; n = 5 resampled runs, fixed session number, varying neuron counts; successful reaching held-out data). Small dots: individual runs. **g:** Mouse forelimb reaching task (adapted from (43)) and recovered 6D latent manifold (first three dimensions shown) trained on DeepLabCut 3D tracking. Points colored by hand position along the reach-to-pull trajectory. **h:** Cross-level decoding. Each node is a level of description of the movement (legend); arrows go from the level used as auxiliary variable to the level decoded from the resulting embedding, and loops show decoding of the level used for training. Line width: decoding R^2^ (%), with the highest values labeled. Full matrix in Supplemental Figure S1d.

We first benchmarked our new sampling scheme on the Neural Latent Benchmark (NLB) datasets. Here, we combined data across various non-human primate limb tasks using velocity as the common label, as the hand position and timing of movements varied across tasks. Specifically, we trained a Unified CEBRA encoder on all six NLB datasets (10), spanning maze, random-target, and bump tasks (Supplemental Figure S1a) and computed behavioral decoding performance for each task, post hoc, from the resulting embeddings. We also compared downstream decoding to POYO, a larger transformer-based multi-session model (39). The unified embedding retained velocity information for every dataset (R^2^ = 0.86–0.95), more than single- or multi-session CEBRA embeddings or POYO (Unified CEBRA higher than CEBRA, CEBRA multi-session and POYO for all datasets; Mann-Whitney U test, 10 runs, Holm-corrected, *p* = 6.5*×* 10^*−*5^ for every test; Figure 1d, Supplemental Table S5).

Notably, Unified CEBRA uses the auxiliary label (here velocity) at inference to select which timepoints of the other sessions are concatenated with the target session. Its scores therefore measure how much task information the shared space retains for the pooled population, not how well behavior can be predicted from one session’s neural activity alone. We benchmark against POYO because it is the closest multi-session model that is also built on behavioral labels: it is trained with behavioral decoding as a supervised loss, building the latent space to specifically target neural decoding. New sessions are likewise added by fitting unit and session embeddings to labeled behavior (i.e., velocity is used as a label). We also compared model fit and stability across runs and found that Unified CEBRA reached a higher goodness-of-fit and a higher cross-run consistency than the CEBRA variants (Supplemental Tables S1, S2). NLB leaderboard results, obtained with methods on a private test split, are listed for context only (Supplemental Table S5).

To evaluate scalability, we applied Unified CEBRA to ninety-nine sessions recorded from three macaques performing center-out (CO; 78 sessions) and random-target reaching (11)(RT; 21 sessions; Supplemental Figure S1b). This then integrated 9,725 neurons into a single embedding (CO: 100.27 *±*9.51, RT: 90.67*±*7.88 neurons per session; Supplemental Figure S1c). Velocity information was highly represented (on successful reaches R^2^ = 94.84*±* 0.11% on CO vs. 95.12% for POYO-MP, and 94.30*±* 0.30% vs. 87.88% on RT; Figure 1e, Supplemental Table S6). Velocity decoding increased steeply with the number of pooled sessions and plateaued beyond 20 sessions (2034.8*±* 231.8 neurons; R^2^ *>* 0.9; Figure 1f). We also note that it only takes 0.3 GPU-hours (GPUs *×* training time) of training, compared to approximately 384 reported for POYO-MP 39 (Supplemental Table S4), thus the similar performance of Unified CEBRA with a computationally much less complex model compared to POYO is notable.

Whereas the non-human primate datasets above use hand velocity, the same movement can be described at several levels of the sensorimotor hierarchy, from hand position and velocity to joint angles, joint torques, muscle forces, muscle lengths and proprioceptive feedback. In our framework, the auxiliary variable plays a role analogous to the design matrix of a generalized linear model (GLM), namely, one can design the auxiliary variables to test a multitude of features, not just one. Importantly, the labels do not impose structure on the embedding. We find that the auxiliary variables unrelated to the neural activity, such as shuffled labels, yield unstructured spaces (Supplemental Table S3). The labels do, however, define which relationships between behavior and neural activity can be tested (as in a GLM setting for single neurons).

We therefore asked how the choice among these descriptive levels affects what the pooled embedding retains. We used two-photon recordings from layer 2/3 excitatory neurons in mouse forelimb somatosensory and motor cortex (22 sessions, 6 mice) during a reach, grasp and pull manipulandum task (43, 60). We derived all six levels of representation using 3D hand and wrist tracking with DeepLabCut and a biome-chanical model of the mouse forelimb (Figure 1g) (43). We trained a unified encoder with each level as the auxiliary variable and measured how much information about the feature each resulting space contained by decoding every level from each embedding space (Figure 1h, Supplemental Figure S1d).

Although the neurons were identical, the extracted spaces differed substantially depending on the auxiliary variable. The spaces conditioned on muscle force and proprioceptive feedback retained the most information about movement between levels (mean decoding R^2^ across target levels: 60.0% and 59.2%), while those conditioned on joint torques retained the least (12.7%). Because an auxiliary variable can only reveal structure that is already present in the activity, these differences reflect how strongly the population relates to each level of description. Choosing auxiliary variables is thus a modeling decision much like choosing regressors in a GLM, i.e., each choice sets which hypotheses can be tested, and comparing across choices shows which descriptions of behavior the neural population most closely reflects.

### The shared latent space is disentangled

Decoding shows that the embedding carries user-defined auxiliary information, but not how that information is organized. A shared space could still mix behavioral variables, a risk specific to pooling sessions with different neurons, so we tested whether the subspace encoding one variable carried little information about another, first in the primate embeddings (reach direction and speed; Supplemental Table S7), and for ease of human interpretability, secondly, we demonstrate this in a well-studied open-source hippocampus dataset 12. In brief, hippocampal CA1 neurons were recorded from four rats running on a 1.6 m linear track (12; Figure 2a). The unified embedding retained more position and direction information than CEBRA and CEBRA multi-session (21) (Figure 2b-c, Supplemental Table S10). It also was consistent across sessions and runs (Supplemental Figure S2a, Supplemental Table S2), with higher goodness of fit (Supplemental Table S1) and position and direction decoding in each individual recording than both CEBRA variants (two-sided Mann-Whitney U test, 10 runs, Holm-corrected, *p* = 6.5*×* 10^*−*5^ for every test; Figure 2c, Supplemental Table S10).

**Figure 2.**
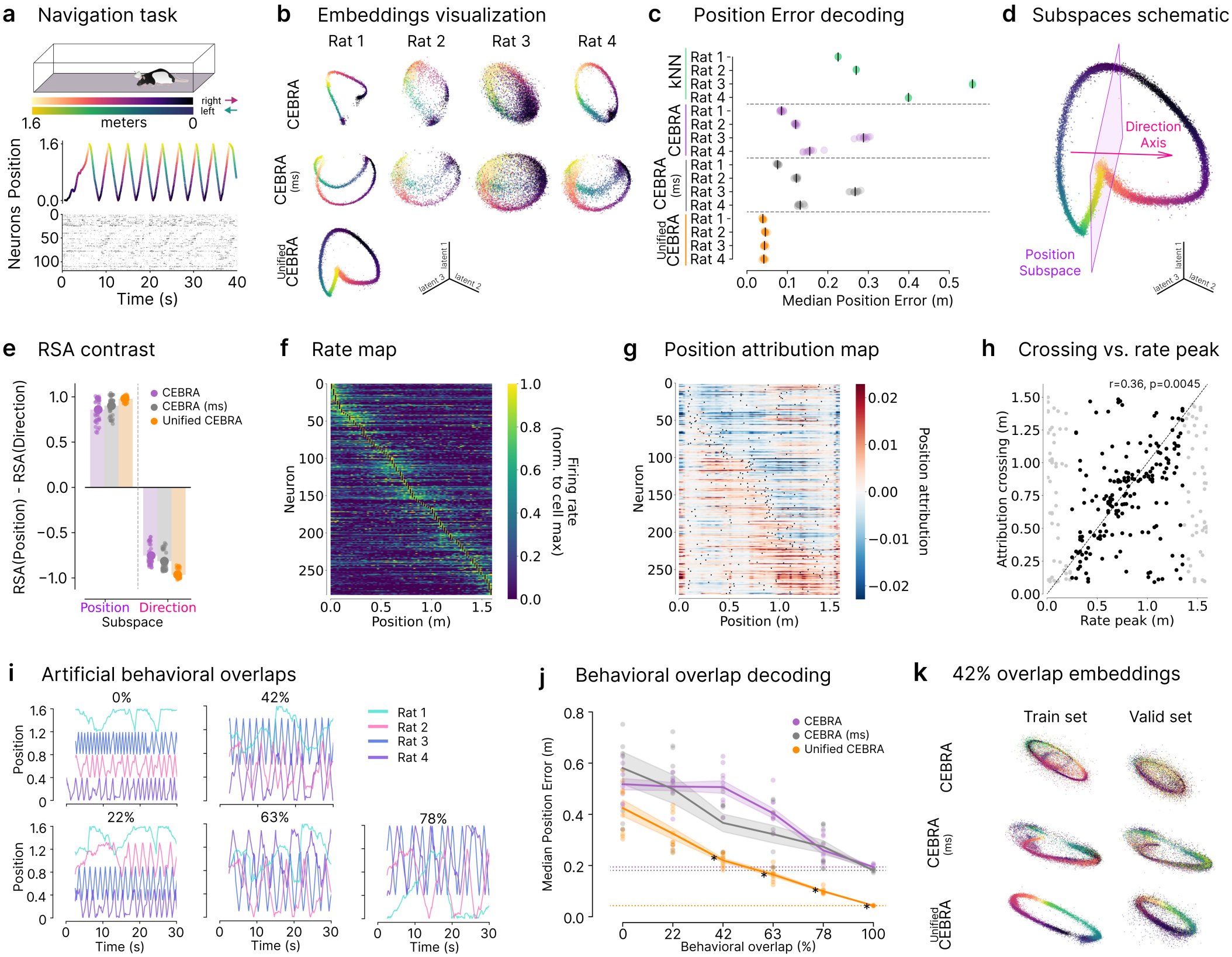
Geometrical disentanglement and robustness to partial behavioral sampling. **a:** Cartoon of the linear track navigation task and neural and auxiliary-label data, adapted from (21); original data are from (12). **b:** Latent embedding visualization (3 out of 32 dimensions). Individual rat embeddings for CEBRA aligned with Procrustes alignment (59), alongside the Unified CEBRA global embedding. Colors matching track position in **a. c:** Median position error (meters; held-out data) from the session-indexed embeddings per rat across runs (n = 10; ticks) and per run (dots). **d:** Schematic of the position subspace and direction axis, drawn over the embedding of a Unified CEBRA model (3 of 32 latent dimensions; colors matching track position in **a**). Projected onto the position subspace, both running directions overlap and position varies; projected onto the direction axis, the two directions separate. The subspaces were estimated in the full embedding (Methods); those shown are illustrative. **e:** Representation disentanglement as ΔRSA = RSA_pos_ *−* RSA_dir_ in the position subspace and on the direction axis. Small dots: individual rats and runs; large dots and bars: mean*±* SEM. **f:** Rate maps of all cells (283 neurons, four rats; training split), each normalized to its maximum and sorted by the position of its peak (black dots). **g:** Position attribution of the same neurons along the track (validation split; mean over five models; same order as f). Black dots: stronger positive-to-negative zero crossings (Methods). **h:** Position of the attribution crossing versus the place field peak for each neuron. Black: neurons scored (rate peak between 0.25 and 1.35 m, n = 179); gray: excluded, near the track ends. Dashed line: identity. r and p: Pearson correlation and empirical p-value against a circular time-shift null (2,000 shifts). **i:** Example artificial position traces for increasing behavioral overlap across sessions. Traces show 30 s of position for each session (rat); in each run, sessions were randomly assigned to a range. **j:** Median position error (meters; held-out uncropped data) from the global embeddings (100% coverage differ from **c**) across runs (n = 10 per condition), as a function of behavioral overlap between recordings during training. Dashed lines: 100% coverage baseline performances. *\**: Unified CEBRA significantly lower than both CEBRA variants at the same coverage, one-sided Mann-Whitney U tests, n = 10, Holm-corrected, *p ≤* 4.1 *×*10^*−*4^). Dots: individual runs; lines: mean*±* SEM. **k:** Example global training and validation sets embeddings for 42% average behavioral overlap across sessions at training (3 out of 32 dimensions; colors matching track position in **a**). CEBRA variants session embeddings aligned with Procrustes alignment (59).

On a linear track, hippocampal place fields are direction-selective (61), so position and running direction can be mixed at the level of single neurons. A useful latent space should nevertheless keep them in separable subspaces, and pooling sessions could plausibly blur them. To measure disentanglement between subspaces, we estimated a position subspace and a direction axis by linear regression of the training embedding on position and running direction (Figure 2d, Supplemental Figure S2b; Methods). On held-out data, we computed representational dissimilarity matrices (RDMs) between conditions (position bin*×* direction) within each subspace (Supplemental Figure S2c), and compared them with the dissimilarities predicted by position and direction. For all three methods, the two variables were separated: the position subspace carried little direction information, and the direction axis carried little position information (double dissociation above the 95th percentile of a circular time-shift null for every rat and run; Figure 2e, Supplemental Table S7). Separation was strongest for Unified CEBRA (*p* = 1.1 *×*10^*−*5^ for all comparisons, two-sided Mann-Whitney test). Pooling sessions, therefore, sharpened the encoding of position and direction.

Disentanglement at the population level does not reveal which neurons drive each latent, a question that matters when the shared space pools sessions with different neurons. We therefore attributed the activity of each neuron to the latents (position, direction) over time (Figure 2f, g, Supplemental Figure 2d; Methods). Neurons’ attributions were mostly structured around their own place field, as one would expect (*r* = 0.36, *p* = 0.0045; Figure 2g-h), indicating that the unified embedding preserves single-cell tuning despite pooling multiple sessions.

### Unified CEBRA allows for partial behavioral coverage

Recording neural activity in freely moving animals means the behavior can be highly variable by design. Such experiments are essential for studying naturalistic activity, but have the problem of non-universal coverage of the behavioral (and possibly neural) space. Methods that aim to find universal encoding embeddings such as Unified CEBRA therefore must tolerate this diversity. While the sensorimotor datasets above already demonstrate different tasks with a common reference (hand velocity) behaviors can be meaningfully combined, we aimed to formally study the overlap tolerance of Unified CE-BRA.

To measure how much behavioral overlap our method tolerates, we simulated that each animal only covered a specific range of space on the track (25–75% of its length), so that it overlapped with the other sessions from 0% to 78%, assigning different subsets to different rats, and evaluated on the full, uncropped held-out data (Figure 2i, Supplemental Table S8). The unified encoder achieved a lower position error overall, and significantly so from 42% (Holm-corrected Mann-Whitney tests, *p≤* 0.002; Figure 2j). With only 63% overlap between sessions, Unified CEBRA reached a lower error than CEBRA and and CEBRA multi-session models trained on the complete track (16.5 vs. 19.4 and 18.1 cm; one-sided Mann-Whitney U test, n = 10, Holm-corrected over 2 comparisons, *p* = 0.009 and 0.014; Figure 2k, Supplemental Figure S2e). A shared manifold can thus be reconstructed even when no single animal samples the entire behavioral space.

We have now established that the behavioral label can be used, much like a GLM design matrix, to both align and query the embedding space. Yet, it is highly important to note that the behavioral label cannot force structure into the latent space. To demonstrate this, we shuffled the neural activity while keeping the label the same: if the label was the primary driver of the embedding, we would be able to decode the behavior; yet this was not possible (Supplemental Table S9). Specifically, shuffling each neuron’s activity in time at inference with labels and matching intact, collapsed hippocampal behavioral decoding (position R^2^ from 0.78 to *−*1.12), showing that label-guided assembly alone does not carry the information.

### Mapping neurons to the latent space

Population-level latent models describe how neural activity as a whole encodes a stimulus, but leave open which neurons carry that information; single-neuron tuning answers the opposite question, with each neuron taken in isolation. Because every neuron enters the same Unified CEBRA encoder, the two can be linked: each neuron receives a signed score for how much it contributes to the population’s readout of a given stimulus feature, on one scale across sessions and animals (Methods, building on 56).

We applied this to 2,924 excitatory (Py) and inhibitory (Sst, Vip, PV) neurons from three visual areas of the Allen Brain Observatory dataset, recorded in 234 sessions from 149 mice viewing a repeated natural movie (13) (Figure 3a-b, Supplemental Table S13). With about 12 neurons per session on average, this means that combining neurons is important to build meaningful embedding models (i.e., prior work showed that 900 neurons and 128 latents best explain the data (21)). As in 21, the auxiliary variable was a pretrained vision-model (DINO) embedding of each frame, and we read out six interpretable movie features from the latent space (motion energy, luminance, vertical and horizontal edges, and low and high spatial frequency), giving each neuron a signed attribution per feature (Figure 3c, Supplemental Figure S3c). The attribution map was reproducible across independently trained models (Supplemental Table S14) and depended on both the trained weights and the auxiliary labels (Supplemental Figure S3d).

**Figure 3.**
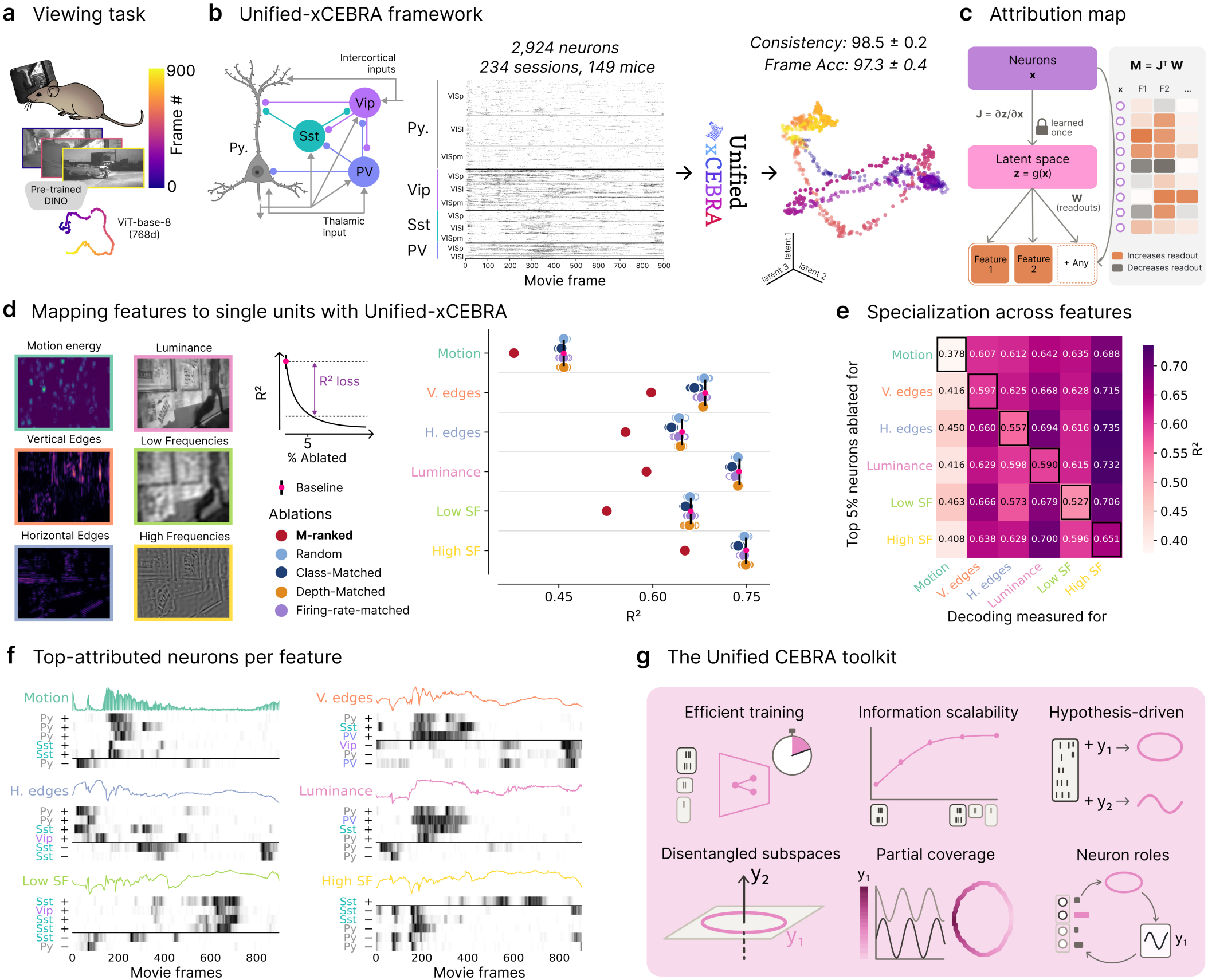
Aunified attribution map identifies feature-specific neurons across recordings. **a:** Mice passively viewed a natural movie (10 repeats); DINO features of the movie frames (visualization: t-SNE; pretrained ViT/8 vision transformer, 768 D feature space) served as the auxiliary variable (adapted from (21)). **b:** Excitatory (Py) and inhibitory (Sst, Vip, PV) neurons from three visual areas (2,924 neurons, 234 sessions, 149 mice) were pooled into a single unified encoder (circuit schematic adapted from (47)). Middle: Activity of all neurons over movie repeat 0 (thick lines), within a type by area (thin lines) and session. Right: shared embedding (3/128 dimensions), with cross-run consistency and frame identification accuracy (kNN; mean*±* SEM, 9 models). **c:** The encoder maps neural activity to the latent space through **J**, and linear readouts **W**, added after training, map it to features. The attribution of neuron *n* to feature *f*, 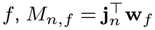, is how far a change in the neuron’s activity moves the latent state along the feature’s readout direction: positive, negative, or near zero depending on their alignment. A rotation of the latent space, as between training runs, leaves the projections unchanged. **d:** Decoding of each movie feature (left; R^2^) after removing 5% of neurons, those ranked highest by the map for that feature (M-ranked) or control sets (random, or matched for cell class and area, imaging depth or firing rate; small circles: individual draws, large circles: mean). Black bar: decoding with all neurons (baseline). Inset: the decrease from baseline after removing a set (R^2^ loss) measures how much decoding depends on those neurons; lower R^2^ means a larger effect. **e:** Decoding R^2^ of each feature (columns) after removing the 5% of neurons ranked highest for each feature (rows). Features differ in baseline decodability, so values should be compared within columns. Solid square: most affected decoded feature (lowest R^2^ in its column). **f:** Responses of the six top M-ranked neurons for each feature (gray), aligned with the feature over the movie (color). **g:** Summary of the properties of Unified CEBRA shown in this study. Training is efficient; the information retained in the embedding scales with pooled recordings; distinct behavioral variables are encoded in separate latent subspaces; recordings can be robustly combined even when their behavioral space does not fully overlap; different auxiliary variables reveal different aspects of the population’s structure; and the same model links neurons to the population representation, and each feature’s readout can be attributed back to neurons across recordings.

The map identified, for each feature, a small set of neurons on which its decoding depended. Removing the 5% of neurons with the largest attributions for a feature, and retraining decoders on the remaining recorded activity, impaired decoding of that feature far more than removing neurons matched for firing rate, imaging depth, or cell class and area (Figure 3d; Supplemental Table S15). The effect was feature-specific: for every feature, removing its own top-ranked neurons reduced its decoding more than removing the neurons ranked for any other feature (Figure 3e). This margin was largest for low spatial frequencies, and smallest for vertical edges. The top-ranked neurons came from every class and area, and their responses tracked the feature across the movie (Figure 3f).

These neurons were largely, but not only, the most strongly tuned: removing the most tuned neurons impaired decoding somewhat less than removing those ranked by the map, for sets of 1–10% of neurons (Supplemental Figure S3e; Supplemental Table S16). Interestingly, the two rankings diverged along cell-type lines. Neurons favored only by tuning were strongly tuned but unreliable across repeats, shared much of their trial-to-trial variability with simultaneously recorded neurons, and were enriched in Vip and PV interneurons; neurons favored only by the map were reliable, shared little variability with their neighbors, and were enriched in Sst and excitatory neurons (Supplemental Table S17).

Because the map keeps the sign of each contribution, classes can also be compared directly. Nine class-feature pairs differed in how often their neurons enhanced the readout, seven in agreement with single-neuron tuning (Supplemental Figure S3f). Sst neurons, for example, more often enhanced the readout of luminance and low spatial frequencies, consistent with their integration of visual input over large regions of space (62). Together, these results are consistent with the encoder weighting neurons by reliability and shared variability as well as by tuning, as population coding theory predicts (63).

## Discussion

We introduced Unified CEBRA, which trains a single contrastive encoder on recordings that share no neurons by assembling label-matched pseudo-populations. Existing multi-session approaches pass each recording through session-specific read-in layers or unit embeddings (38, 39, 42). In Unified CEBRA, every neuron enters the same encoder, so recordings reinforce one another. The pooled embedding retained more task information than single- and multi-session CEBRA embeddings, matched or exceeded the multi-session transformer POYO, separated behavioral variables more cleanly, and tolerated partial behavioral overlap, while training in minutes on a single GPU. New recordings can be added by training only their input channels, leaving the pretrained encoder unchanged (Supplemental Table S10).

The auxiliary variable can be a repeated stimulus or, in principle, a discrete task event such as trial start; kinematic or biomechanical variables; or learned representations such as DINO features. Thus, it can be a ‘harder’ constraint (exact 2D position), or a task-level feature that is arguably let restrictive on the learned latent space. Importantly, it sets which hypothesis is tested without imposing structure, as our shuffle controls show. Because the auxiliary variable is required at inference, where it selects the timepoints pooled from other sessions, Unified CEBRA is designed for offline analysis of experiments in which that variable is known, not for decoding unknown behavior or for online use. The unified embedding assumes that sessions share a latent state at matched values of the auxiliary variable; differences in internal state or strategy across animals violate this assumption to some degree, and cross-session consistency measures how closely it holds. Because sessions are concatenated, the input dimension grows with the total number of neurons, which will eventually limit scaling to very large collections.

Pooling also yields a run-invariant mapping from neurons, through a shared non-linear and identifiable latent space, to features. Whereas xCEBRA (56, 57) attributes latent factors to neurons by inverting the encoder, chaining the encoder’s Jacobian with linear feature readouts attributes the readout of each feature to neurons across sessions, placing every neuron on one signed scale that is invariant to the linear indeterminacy of contrastive learning. Attention weights are not identifiable (50, 52), readouts of predictive models change coordinates between runs (44–46).Our map describes each neuron’s role in the population readout, bridging population codes and single-cell analysis. In mouse visual cortex (Allen Dataset), with about 12 neurons per session on average, it identified feature-specific neurons that were not only strongly tuned but also more reliable and, where it departed from tuning, less coupled to their neighbors, consistent with population coding theory (63). Whether attribution-selected ensembles are more behaviorally effective than those selected by visual sensitivity (64) remains to be tested *in vivo*. Additionally, large multi-region datasets with rich behavioral labels (65) and longitudinal recordings of the same (or different) neurons across learning (66) are natural next applications.

## Code Availability

All code is available under an Apache 2.0 license at: https://github.com/AdaptiveMotorControlLab/CEBRA version 0.6.1 onward.

## Dataset Availability

- Neural Latent Benchmark dataset: https://neurallatents.github.io/datasets.
- Perich dataset: https://dandiarchive.org/dandiset/000688 and we used the pre-processing proposed in (39).
- Hippocampus dataset: https://crcns.org/data-sets/hc/hc-11/about-hc-11 and we used the preprocessed data from https://figshare.com/s/60adb075234c2cc51fa3?file=36869049 (21).
- 2P visual coding (Allen Institute) dataset: https://github.com/zivlab/visual_drift/tree/main/data and we used the preprocessed data from https://figshare.com/s/60adb075234c2cc51fa3?file=36869049 (21).
- M1/S1 dataset: data from (43).

## Acknowledgments

We acknowledge our funding: SNSF Grant 320030-227871 and TMSGI3_226525 (MWM), The Simons Collaboration on Ecological Neuroscience (SFI-AN-NC-SCN-00007276; MWM), and EPFL for funding. MWM is the Bertarelli Foundation Chair of Integrative Neuroscience.

## Author Contributions Statement

Conceptualization: M.W.M., C.B., St.S.; Software: C.B., St.S., H.M., M.W.M.; Investigation: C.B., H.M., St.S.; Writing-Original Draft: M.W.M., C.B., St.S., H.M.; Supervision & Funding Acquisition: M.W.M.

## Declaration of interests

EPFL has a patent relating to the use of CEBRA for brain-machine interfaces (US Patent No. US12499131B2), where St.S. and M.W.M. are listed as the co-inventors. The authors declare no other competing interests.

## Methods

### Mathematical details of Unified CEBRA

#### Initial Intuition

We assume that neural activity recorded in different sessions reflects a shared underlying latent structure tied to observable variables such as time within a task, behavior, or trial events, even though the identity and number of recorded neurons vary across sessions. Sessions are considered alignable when these auxiliary variables are defined in a comparable way, making it possible to relate neural activity to the same behavioral or task states. To align sessions, we use auxiliary variables as anchors: for a given auxiliary state (e.g., a specific moment in the task or behavioral configuration), each session contributes the neural activity recorded at its closest matching value. Combining these samples forms a pseudo-population representation of the same latent state across sessions. We then train a single encoder using a contrastive objective that encourages representations corresponding to similar auxiliary states to be close, while separating representations associated with distinct states, yielding a session-invariant neural population representation.

#### Mathematical formulation

Mathematical formulation. Suppose we have *M* sessions, where the *i*-th session contains neural data *X*_*i*_ and auxiliary variables *Y*_*i*_. Formally, 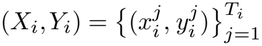, denotes the data recorded at time step *j* for session *i*. Here, 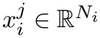 is the population activity of *N*_*i*_ neurons in the session at time *j*, 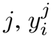 is a corresponding auxiliary variable (e.g., time, a behavioral, or stimulus label), and *T*_*i*_ the total number of time steps in the session. Let us denote with 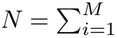 *N*_*i*_ the total number of neurons across all sessions.

Our goal is to construct a single encoder *f* : ℝ^*N*^ ↦ ℝ^*D*^ capable of processing data from all *M* sessions, even though each session has different dimensionalities *N*_*i*_, and mapping the shared activity onto a lower dimensional latent space with dimensionality *D*. Because a typical encoder requires a fixed input size, we develop a multi-session alignment strategy that maps data from all sessions into a shared latent space. Specifically, we concatenate samples from all sessions into one fixed-dimensional vector, using the auxiliary variables *Y*_*i*_ to align samples with similar behavioral states.

#### Contrastive learning

Let *f* be a neural network encoder parameterized by the weights *θ*. Contrastive learning aims to learn an encoder *f*_θ_ that maps samples into an embedding space where similar samples (positive pairs) lie closer, and dissimilar samples (negative pairs) lie farther apart. Formally, let *x* be a reference sample, (*x, x*+) a positive pair, and (*x, x*^−^) a negative pair. We train *f*_θ_ to maximize *ϕ*(*f*_θ_ (*x*), *f*_θ_ (*x*+)) and minimize *ϕ*(*f*_θ_ (*x*), *f*_θ_ (*x*^−^)), where *ϕ* is a similarity measure (here, we use the dot product between normalized vectors). Specifically, we leverage the following InfoNCE loss function:

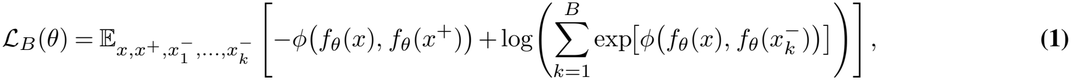

where *B* is the batch size. By minimizing ℒ_*B*_(*θ*), the encoder learns to embed samples with similar auxiliary labels closer together, while pushing away those that differ.

#### Forming reference samples and their pairs from multiple sessions

We propose a new sampling scheme for reference, positive and negative samples which naturally aligns neural activity across sessions, enabling a single encoder to handle multi-session neural data from diverse sources.

##### Reference Samples

At each training step, we randomly permute the session labels (1, …, *M*) and denote this permutation by the tuple *S*. We denote the *reference session* as *S*_1_. From this session, we randomly sample a reference time step *r*_1_ corresponding to the sample 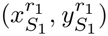. For every other session *S*_2_, …, *S*_*M*_, we find the timestep whose auxiliary variables 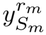 is closest to 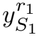, using a chosen distance metric:

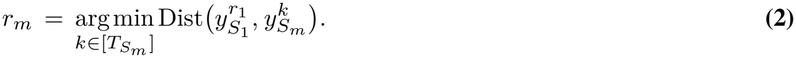

We chose Dist to be the squared Euclidean distance and *k* is selected from the possible indices 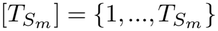 of the whole session *S*_*m*_. The distance function can be adapted to the data modality of the auxiliary variable. If several timepoints are equally close, one is selected uniformly at random among them, whereas the original CEBRA implementation selects the first tied timepoint (21).

Once these matches are identified, we concatenate their neural activity vectors into a single fixed-dimensional reference:

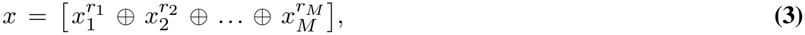

where *⊕* denotes concatenation along the neuron dimension, resulting in a vector of size 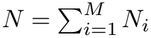, ordered by session identity.

##### Positive Samples

For each reference sample index *r*_*m*_, we construct a positive sample from the same session *S*_*m*_, by selecting a time step at which the auxiliary variable is sufficiently close. Similarly to Schneider et al. (21), we first compute the set of empirical differences 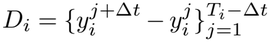 to *r*_*m*_ for each session *i*, where the “time offset” Δ*t ∈* ℤ + is a user-defined fixed hyperparameter that defines the temporal scale for auxiliary-space variability. Uniformly sampling a difference Δ*y*_*i*_ from each *D*_*i*_ approximates sampling from the empirical distribution of changes in the auxiliary variable. Per session *S*_*m*_, we determine the closest match:

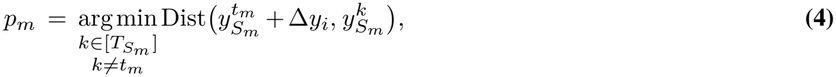

where Dist is the same auxiliary-space distance used for reference alignment. Finally, we construct the overall positive sample by concatenating across all ordered sessions:

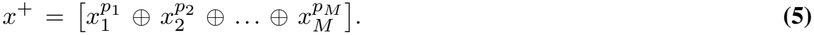

By introducing this localized random selection, we effectively diversify the positive sampling strategy, which has been shown to enhance representation learning in contrastive objectives (67, 68).

##### Negative Samples

For each negative pair, we randomly sample a data point from the reference session *S*_1_, denoted 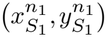. For each remaining session *S*_*m*_ (*m ≠* 1), we select the associated sample 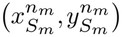 such that 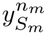 is close to 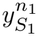analogous to Eq. 2. We then concatenate these points across ordered sessions, yielding

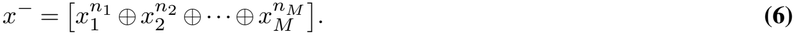

For each reference sample, we create multiple negative pairs and the number of negative samples is a hyperparameter of the algorithm which matches the batch size.

#### Label-guided inference

To infer our unified representation, we used a sampling procedure similar to that used at training. For a given target session *T*, each data point 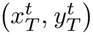 at timepoint *t* is matched to the closest point in each other session *j* in terms of the squared Euclidean distance in the auxiliary space: 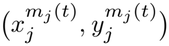 . The encoder input is the concatenation:

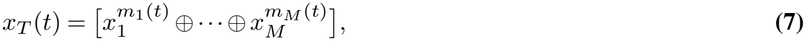

where *m*_*T*_ (*t*) = *t*. The resulting trajectory *z*_*T*_ (*t*) is thus an embedding of the pooled population, indexed by the behavioral timeline of session *T* . Auxiliary labels are never passed to the encoder; they only determine which timepoints of each session are combined, and at test time matched timepoints are drawn from the held-out split of every session.

Repeating this procedure for every session *T∈{* 1,…, *M}* yields *M* session-indexed trajectories *z*_1_, …, *z*_*M*_, all produced by the same encoder and expressed in a single latent space. Concatenating them in time gives what we refer to as the *global embedding*: a representation of the pooled population that spans the behavioral distribution of all sessions. For CEBRA variants, a comparable global embedding is obtained by aligning the session embeddings with Procrustes alignment (59) before concatenation.

#### Neural data masking

Following masked models of neural activity, which mask neurons and timesteps during training to discourage reliance on any single neuron or timebin (41, 42, 69–73), we applied stochastic input masking during training on all but the regularized Unified CEBRA models (all datasets but the 2P Allen dataset). For each reference (*x*), positive (*x*^+^), and negative (*x*^*−*^) sample within a batch, we independently sampled a masking scheme and a masking ratio. We used combined neuron and time masking with a flexible ratio (0.3–0.95, step 0.05), as in masked models of neural activity (42, 71).

#### Cross-subject adaptation

We evaluated cross-subject adaptation under this masked pretraining scheme using a leave-one-out framework across recordings. After pretraining on three subjects out of four, all pretrained layers were frozen, and only the input channel weights for the held-out subject were trained for 100 steps (50-fold fewer than training from scratch), matching *de novo* performance (hippocampus dataset; Supplemental Table S10).

#### Neural shuffle controls

To test whether decoded information emerges from neural activity patterns or from label-guided sampling, activity was permuted independently in time for each neuron, while leaving the labels and the matching procedure intact. This preserved firing statistics but removed the relationship to auxiliary variables and correlations between neurons. We applied this shuffle in two settings:

- *At inference*: the encoder was trained on intact data and the held-out data were shuffled before label-guided assembly: if matching by label alone carries the decoded information, decoding should persist despite the shuffle.
- *At training*: the encoder was trained on shuffled data and evaluated on intact held-out data, testing whether the contrastive supervision imposes the label structure on the latent space independently of the neural data, as in (21).

In each setting, we shuffled either all sessions or all sessions except the target session (tested for all), the latter quantifying the contribution of the other sessions. All conditions used the hyperparameters of the intact model, and the same kNN decoder (k = 36) fit on the training embedding and evaluated on held-out data.

### Additional Evaluation Metrics

**Goodness-of-fit (GoF)**. To measure how well encoders fit the data, we used the goodness-of-fit (GoF) (21, 43), in bits:

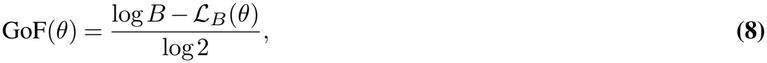

where ℒ_*B*_(*θ*) is the InfoNCE loss (Eq. 1) of a converged model *θ* and *B* the batch size. For CEBRA multi-session, *B* is the cumulative batch size, i.e., the batch size per session times the number of sessions. A GoF of 0 bits denotes chance level, and higher values indicate a better fit. Because its scale depends on batch size, temperature and the behavioral structure of each dataset, GoF is used only to compare models within a dataset. Since input masking lowers the GoF computed during training, when reported, we report GoF on held-out data for all models (NLB, Perich and hippocampus datasets; Supplemental Table S1). For the visual cortex dataset, whose model was trained without masking, we also report the GoF during training, to show its stability across runs.

#### Consistency between embeddings

Consistency between two embeddings is measured as the R^2^ score of a linear regression (including an intercept) predicting one embedding (target) from the other (source). We report this as a matrix of R^2^ scores for each pair. Note that these matrices are not symmetric, as the linear regression is directionally asymmetric and yields two values per pair. We distinguish two types of consistency. Cross-run consistency compares models trained on the same data with the same parameters, and reflects the stability of training across random seeds: a model with low cross-run consistency is unreliable regardless of its other properties, so we use it to select the best hyperparameters for each dataset. Cross-session consistency compares embeddings from the same trained model type applied to different recordings. Because our Unified CEBRA encoder projects all sessions into a shared latent space, we expect cross-session consistency to align with cross-run consistency if the model has learned a truly session-invariant representation. We use cross-session consistency as a control, to confirm that Unified CEBRA achieves higher cross-session consistency than the CEBRA baselines.

#### Separation of behavioral variables in subspaces

Because embeddings trained with cosine similarity are identifiable only up to an orthogonal transformation, individual latent dimensions have no fixed orientation, and we therefore measured how behavioral variables were encoded in subspaces of the embedding using representational similarity analysis (RSA). We describe the analysis for the hippocampus dataset (position and running direction), and its adaptation to the primate dataset below.

For each rat and run, a position subspace and a direction axis were fitted on the training data by linear regression of the embedding on position and running direction. Each timepoint *t* has an embedding **z**(*t*) ∈ ℝ*D*, a position bin *b*(*t*) ∈ {1,…, 8} (0.2-m bins along the track) and a running direction *d*(*t*) ∈ {0, 1}, and we modeled

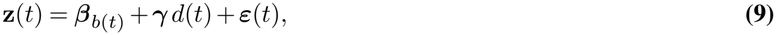

where *β*_*k*_ ∈ ℝ^*D*^ is the effect of position bin *k* and **γ** ∈ ℝ^*D*^ the effect of running direction. Both were estimated jointly by least squares (one-hot regressors for the bins, one regressor for direction), so that each effect is controlled for the other. The direction axis is the unit vector of the direction effect **u** = **γ**/‖**γ**‖. The position subspace was obtained by principal component analysis of the mean-centered bin effects. We formed 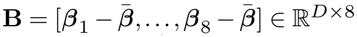, computed its singular value decomposition **B** = **UΣV**^⊤^, and defined the position subspace **P** ∈ ℝ^*D*⨯*?*^ as the first *k* columns of **U**, with *k* the smallest number of components whose cumulative variance reached 90% (*k* = 2 for all 40 unified and multi-session rat × run units; *k* = 1 or 2 for single-session units, 19 and 21 of 40). On held-out data, the embedding was averaged within each of the 16 conditions (8 position bins × 2 directions), giving condition means **m**_*k*_,_*d*_, which were projected onto each subspace, **p**_*k*_,_*d*_ = **P**^⊤^**m**_*k*_,_*d*_ and *q*_*k*_,_*d*_ = **u**^⊤^**m**_*k*_,_*d*_.

For each subspace, the representational dissimilarity matrix (RDM) contains the Euclidean distances between projected condition means 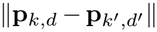 and 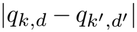. It was compared with a position RDM, 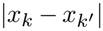 with *x*_*k*_ the center of bin *k*, and a direction RDM, equal to 1 for conditions with different directions and 0 otherwise, by Spearman correlation of their upper-triangular elements. For each subspace, we computed the RSA contrast ΔRSA = RSA_pos_™ RSA_di?_, and defined the double dissociation as

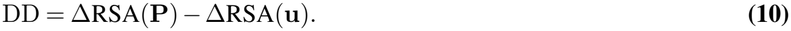

DD reaches 2 when the position subspace reflects only position and the direction axis only direction.

For each rat*×* run unit, the double dissociation was compared with a null distribution obtained from 2,000 circular time shifts of the labels, applied before the subspaces were fitted (one-sided empirical p-value). Models were compared with the run as the unit, after averaging over rats (two-sided Mann-Whitney test, 10 runs per model).

For the primate reaching datasets (NLB and Perich), the same analysis was applied with reach direction and speed, both derived from hand velocity, the auxiliary variable. Reach direction (the angle of the velocity vector) was divided into 8 angular bins, and speed (its norm) into 4 bins, with bin edges computed on the training split of each session; the slowest 10% of samples were excluded, since direction is undefined at rest. Both subspaces were obtained as for position: a joint linear model with direction and speed bin indicators was fit, and each subspace was spanned by the principal axes of its mean-centered bin effects that explain 90% of their variance. The direction RDM used the circular distance between bin centers, and the speed RDM the absolute difference in speed-bin rank. RSA was computed on the condition means of the direction speed grid (conditions with at least 20 samples), and the double dissociation defined as above, with the direction subspace in place of the position subspace and the speed subspace in place of the direction axis. Units were runs for NLB (10 runs, averaging the 6 sessions within each run) and training seeds for Perich (5 seeds, averaging the 99 sessions within each seed); nulls and model comparisons were computed as above.

### Regularized unified latent space and neurons contribution

#### Regularized unified encoder

We adapted xCEBRA (56) to the Unified CEBRA framework, using the behavior-conditioned contrastive objective. We also kept its Jacobian regularization (RegCL) rather than the neuron and timestep masking used elsewhere in this work. Among encoders that fit the data equally well, the regularization selects the one with the smallest input sensitivities, which makes the distribution of attributions among correlated neurons unique. The regularization weight was ramped linearly from 0, starting at one quarter of training and reaching its target value at one half, as in (56). The target weight was selected by grid search to jointly maximize cross-run consistency and minimize the final training loss (mean of the last 10 iterations), and each run then was assigned a weight sampled uniformly around this value, so that reliability across also reflects robustness to the regularized strength.

#### Neuron-to-feature attribution map M

To resolve the functional contributions of individual neurons to specific movie features, we use the Neuron Gradient method (74) as in Schneider et al. (56) to compute the encoder’s local Jacobian and composed it with a post-hoc linear decoder. Let **x** ∈ ℝ^*N*^ denote the population neural activity, **z** = *f*_θ_(**x**) ∈ ℝ^*D*^ the latent embedding generated by the unified encoder *f*_θ_(·) : ℝ^*N*^ → ℝ^*D*^ parametrized by *θ*, and **ŷ** = **W**^⊤^**z** ∈ ℝ^*F*^ the *F* continuous movie features estimated by a linear ridge decoder with weights **W** ∈ ℝ^*D*⨯*F*^. Ridge decoders (*α* = 9) were fit with an intercept (unused by the map) from latents to standardized movie features on the nine runs across folds. Because the decoder is linear, all non-linearity comes from the encoder. The local rate of change of decoded features with respect to input neurons is ∂**ŷ**/∂**x** = **W**^⊤^**J** ∈ ℝ^*F*⨯*N*^, where **J** = ∂**z**/∂**x** ∈ ℝ^*D*⨯*N*^ is the encoder Jacobian. To structure attributions with neurons along rows and features along columns (ℝ^*N*⨯*F*^), we define the sensitivity matrix **M** as its transpose:

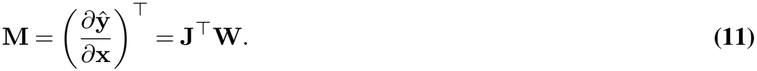

Because the encoder takes as input a window of *K* samples around time *t* (its temporal receptive field), **J**(*t*) was computed with respect to that window and averaged over the window’s samples. We computed the attribution map **M** of each model, the average of **M**(*t*) over the *T* timepoints of the data, equivalently obtained by averaging **J**(*t*) before multiplying by **W**, since **M**(*t*) is linear in **J**(*t*). Each entry is then the dot product 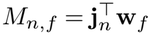 between the sensitivity of the latent state to neuron *n* (**j**_*n*_, the *n*-th column of the time-averaged Jacobian) and the readout direction of feature *f* (**w**_*f*_, the *f*-th column of **W**). The time-resolved maps **M**(*t*) were also used to study how neurons’ contributions vary over time.

Contrastive embeddings are identifiable only up to an invertible linear transformation **z**′ = **Tz**, which is orthogonal for normalized embeddings with cosine similarity (54). Under this reparameterization the encoder Jacobian transforms by the chain rule as **J**′ = **TJ**. For the ridge readout used here, stacking the latents and features of the training timepoints as the rows of **Z** and **Y**, the least-squares solution is **W** = (**Z**^⊤^**Z** + *α***I**)^−1^**Z**^⊤^**Y**, and fitting it on **Z**′ = **ZT**^⊤^ gives **W**′ = **T**^−⊤^ (**Z**^⊤^**Z** + *α*(**T**^⊤^**T**)^−1^)^−1^**Z**^⊤^**Y**, which equals **T**^−⊤^**W** when **T** is orthogonal, as expected for normalized embeddings. The composed forward attribution **M** is therefore invariant across runs:

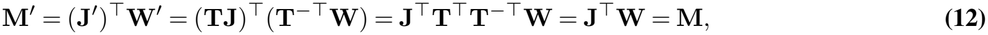

With the ridge readout used here, this holds exactly when **T** is orthogonal, as expected for normalized embeddings, and approximately otherwise.

Compared to the Neuron Gradient method, the Inverted Neuron Gradient method, also presented and validated in Schneider et al. (56), instead attributes the data-generation process, which is approximated by inverting the learned encoder. As a consequence, its attribution uses the pseudo-inverse of the Jacobian. Through the regression approach outlined above, we are interested in the inverse direction, i.e. how each neuron contributes to the readout of a feature. We therefore use the forward Jacobian of the encoder, chained with the readout, as in gradient-based attribution methods.

#### Model reliability and architectural sanity checks

We verified the statistical reliability and architectural dependence of **M** through three complementary controls (75).

- *Pairwise reliability:* For each pair of runs (*r, r*^*′*^) and each feature *f*, we computed the Spearman rank correlation across all *N* neurons between 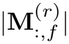 and 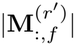, and averaged it over pairs and the *F* = 6 features. At the class level, we computed the Pearson correlation between the directional biases Δ_*c*_(*f*) (Eq. 14, below) of the two runs, across the 24 class*×* feature entries.
- *Cascading parameter randomization:* For each run *r*, we randomized the encoder layers cumulatively, from the output layer backward. At each step, we refit the linear decoder on the perturbed latents, recomputed the map, and computed the Spearman rank correlation of its |**M**_:,*f*|_ with the trained map of the same run, averaged over features (3 runs). As a chance baseline, we permuted neuron identities (rows of **M**^(*r*)^; 200 permutations) for each run, and averaged the resulting null over runs.
- *Auxiliary-Label Shuffling:* We trained three runs with the auxiliary variables (DINO features) permuted in time independently for each session, keeping all hyperparameters identical. Linear decoders were then refit to predict the (unshuffled) movie features from each shuffled run’s embedding, computed with the sessions aligned on the true movie frames, and cross-model attribution reliability was evaluated across all shuffled pairs (3 pairs; see above).

#### Targeted *in silico* ablation assay

To test whether top M-ranked neurons are needed to decode visual features from recorded activity, we implemented an *in silico* targeted ablation assay. Linear ridge decoders were fit directly on the raw calcium traces of the neural population stacked across recordings to predict visual features, bypassing the latent space entirely. Decoders were fit on the training repeats of the run and scored on the held-out repeat, with the regularizer *α∈* [10^0^, 10^7^] optimized via exact leave-one-out cross-validation (`sklearn`).

For each feature *f*, we ranked neurons by their attribution magnitude for that feature, 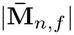, averaged over the three runs of each cross-validation fold, so that both enhancing and suppressing neurons were targeted. We removed the top 5%, refit the decoder on the remaining neurons, and scored it on that fold’s held-out repeat; scores were then averaged over folds. The effect of removing a set of neurons is its R^2^ loss, the decrease in held-out decoding relative to the full population:

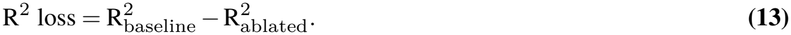

The attribution-ranked set was compared with size-matched control sets (10 draws each):

- *Uniform random:* neurons drawn uniformly at random from the population.
- *Class-matched:* neurons drawn to match the cell class and area composition of the ranked set, ruling out effects of cell-class abundance.
- *Depth-matched:* neurons drawn to match the imaging-depth composition of the ranked set, ruling out effects of recording depth.
- *Firing-rate-matched:* neurons drawn to match the ranked set across 10 deciles of baseline log event rate, borrowing from adjacent deciles when a decile had too few neurons, ruling out effects of activity level.

To test whether effects were specific to the targeted feature, we also investigated the R^2^ loss of each ranked set on the decoding of the other five features. Feature specificity was quantified by assessing diagonal dominance.

To determine how many neurons were needed for an effect, set sizes were varied over *k∈ {*3, 6, 15, 29, 58, 146, 292, 585} neurons (0.1, 0.2, 0.5, 1, 2, 5, 10, and 20%). For each feature, set size and control type, we report the number of control draws whose R^2^ was lower than that of the attribution-ranked set, and the number of features for which the ranked set produced the larger loss.

#### Group-level directional bias

To determine whether a class *c* systematically differs from the others in increasing or decreasing the readout of feature *f*, we computed its directional bias Δ_*c*_(*f*): the fraction of its neurons with a positive attribution, relative to the unweighted mean of that fraction across the *K* classes,

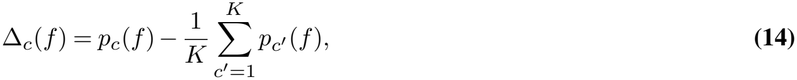

where *p*_*c*_(*f*) is the fraction of neurons of class *c* with 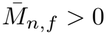. Because this analysis uses only the map and no held-out data, 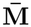 was averaged over all nine runs. Because neurons from the same session are not independent, *p*_*c*_(*f*) was estimated with a logistic mixed model, 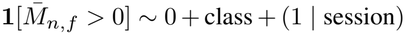, fitted separately for each feature, and reported as the population-averaged fraction (integrating over the session random effect).

Significance of each class’s contrast was assessed by parametric bootstrap (1,000 datasets simulated from the fitted model and refitted; two-sided *p*-values), with Benjamini-Hochberg correction across the 24 class–feature tests.

#### Marginal tuning

As a model-free reference for the attribution map, we computed for each neuron *n* and feature *f* its marginal tuning *r*_*n,f*_, the Pearson correlation at zero lag between the neuron’s trace averaged over the fold’s training repeats and the z-scored feature. Marginal tuning was used as an alternative ranking for the ablation, removing the neurons with the largest |*r*_*n,f*|_ under the same procedure as for 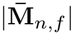, for class-level directionality, replacing the sign of 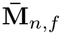 by the sign of *r*_*n,f*_ in the same mixed models.

To quantify agreement between the two rankings, we computed, for each feature, the Spearman correlation across neurons between 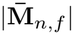 and 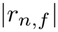 . We then compared the neurons selected by each ranking. For each feature, we took the top 5% of neurons by attribution magnitude and by marginal tuning, and split them into neurons selected by both rankings, by the map only, and by tuning only. Selection used only data independent of the held-out repeat (the models and training repeats of each cross-validation fold), and all results were pooled over folds. For each neuron, we computed two measure from its binary event train. Stimulus reliability was the Pearson correlation between its response on the held-out repeat and its mean response over the training repeats. Within-session coupling was the Pearson correlation, over the training repeats, between its deviation from that mean response and the summed deviations of the other neurons recorded in the same session (sessions with at least three neurons). We compared map-only and tuning-only neurons on each measure and on their cell-type composition (the fraction of each cell type in each group). For each comparison, we computed the difference between the group means within each feature and fold, and averaged it over features and folds. Significance was assessed with a two-sided permutation test that shuffled group labels between map-only and tuning-only neurons within each feature and fold (2,000 shuffles).

### Baseline Models

#### CEBRA Models

We used CEBRA (21), more specifically CEBRA-Behavior in its single-session and multi-session modes, as both a baseline method and a code base to develop our new unifying sampling scheme. In the single-session setting, the sampling procedure consists of distinguishing the correct positive sample from multiple negative samples given a uniformly-drawn reference sample, in one session. In the multi-session setting, it is similar except that the positive sample may originate from a different session provided it matches the behavioral variables of the reference sample and negative samples are drawn across sessions. Then each session is projected back into its own encoder, with its specific neuron count as the input size.

#### POYO Models

POYO (Pre-training On manY neurOns; 39) is a PerceiverIO-based model for multi-session neural decoding. Each spike is represented as a token combining the identity of the unit that emitted it and its timestamp. These tokens are compressed into a fixed set of latent tokens by cross-attention, processed by self-attention layers, and behavior is decoded from output query tokens, one per requested output time, that cross-attend to the latents. Units and sessions are represented by learnable embeddings, stored in a vocabulary initialized from the training data and extensible to new recordings.

On the NLB dataset (Supplemental Table S5), we trained POYO ourselves with the official implementation (https://github.com/neuro-galaxy/torch_brain, commit 3620c46) and the configuration provided for this dataset (pei_pandarinath_nlb_2021 brainset pipeline), on the same splits and with the same evaluation (public validation split; 10 runs) as the other models. The default context window of 1 s exceeds the length of the NLB trials (about 700 ms), and the sampler discards trials shorter than the window, so we reduced it to 0.5 s, keeping all other hyperparameters at the authors’ values. On the Perich dataset, we report the results of POYO-MP, a multi-session POYO model trained on the same 99 sessions from Azabou et al. (39), with the same validation splits and scoring during reaching.

#### NLB Models

On the NLB, we reported the existing top solutions that were submitted to the leaderboard. We only reported them for context only as they were evaluated on the private evaluation split, contrary to the models we trained locally and evaluated on the public validation split.

#### Neural kNN Models

As a baseline decoding control, we evaluated a kNN (k-Nearest Neighbors) model on the original neural high-dimensional space for the NLB datasets. We performed a small grid search to optimize the number of neighbors (*k*) from [1, 2, 5, 8, 16, 24, 36, 48, 64], trained one model per session (as input size differs) and reported the decoding performances.

### Benchmark Datasets

We benchmarked Unified CEBRA on three different types of tasks, widely common in neuroscience, namely motor, navigation, and visual tasks, against existing methods. Across all datasets, encoders were trained using AdamW (weight decay = 0.01) with a 10% linear warm-up followed by cosine decay. Hyperparameters (learning rate, temperature, batch size, receptive field, and embedding dimension) were selected via Ray Tune or Weights & Biases Bayesian sweeps by maximizing cross-run consistency (summary in Supplemental Table S11). Below, we also provide additional specific information.

#### Neural Latent Benchmark

The datasets in the Neural Latent Benchmark (NLB) comprise three different rhesus macaques motor tasks where they make reaching movements in a virtual maze: a ‘Maze’ task, a ‘Bump’ task and a ‘RTT’ task (Random Target Task). Neural activity while the monkeys perform the tasks was recorded in the motor cortex (M1) and dorsal premotor cortex (PMd) (MC) and the somatosensory cortex (Area2) using an 96-electrode Utah array, with Δ*t* = 5 ms (10). For the MC-Maze task, there are four settings: MC-Maze (“Standard”), MC-Maze-s, -m, or -l. The MC-Maze scaled datasets are from separate sessions and differ from the ‘Standard’ one by the number of trials. With MC-RTT, a random target reaching task (98 neurons) and Area2-Bump, a limb perturbation task (49 neurons), this is a total of 6 sessions and 627 neurons.For all tasks the x- and y-velocity average is the behavioral output to decode on the benchmark. We used both x- and y-velocity as the auxiliary variables for training. Smoothing with a Savitzky-Golay filter, window length=9, polyorder=1 of the embeddings was applied before using them for behavior decoding. More dataset details can be found in Pei et al. (10) and https://neurallatents.github.io/datasets.

We trained the CEBRA multi-session implementation (21) and Unified CEBRA on the datasets described above (training split only) so that each dataset is considered as a session in our setup. Single-session CEBRA models (21) were trained on each dataset individually. The x- and y-velocity were normalized (z-scored) per task and used as the auxiliary variables. Following encoder training, we trained a ridge regression model to decode the x- and y-velocity of the hand from the training data output embeddings, and evaluate on the held-out data. We obtain the R^2^ scores for each and report the average velocity R^2^ scores. We also report the held-out goodness-of-fit of the CEBRA baselines and Unified CEBRA models.

#### Perich Dataset

The Perich et al. (11) dataset consists of 112 sessions recorded in four rhesus macaques monkeys performing either a center-out (CO) task, or a continuous random-target (RT) acquisition task. Following Azabou et al. (39), we excluded monkey T (12 sessions; 6 CO sessions, 6 RT) and one CO session from monkey C from the training set and only considered the data from monkeys C, M, and J for a total of 99 sessions (78 CO, 21 RT sessions). We split the data randomly by trial for each session, so that we keep 70% of the trials for training and evaluate on the 30% held-out trials.

Neural activity was recorded from chronically implanted Utah microelectrode arrays in the primary motor cortex (M1) or dorsal premotor cortex (PMd). On average, the sessions contained respectively 100.27 *±*9.51 for CO and 90.67*±* 7.88 neurons for RT sessions, for respectively 7,821 and 1,904 neurons and a total of 9,725 neurons. The data contains spiking activity, manually spike-sorted, cursor position and velocity in 2D, and other task-related metadata. The Perich dataset is available at https://dandiarchive.org/dandiset/000688 and we used the pre-processing proposed in Azabou et al. (39).

The Unified CEBRA encoder was trained across all sessions on the train set for 5 runs using the x- and y-velocity as the auxiliary variables. The embedding was smoothed with a first-order Savitzky-Golay filter with window length 19 before behavior prediction. As in Azabou et al. (39), we computed the velocity R^2^ as the average of the x- and y-velocity R^2^ for each session on the held-out trials, evaluated on the “active” part of the successful trials only (CO: “Reach” phase, RT: “Random”). We do not train the decoder end-to-end with the encoder, only post hoc decoding by training a 2-layer MLP decoder (hidden dimension 32, GeLU activation function, 20 epochs, learning rate 0.001) on the training set latent embeddings and evaluating performances on the valid set. We also reported the cross-run consistency averaged across sessions. We compared Unified CEBRA to POYO, using the results directly provided by (39) for their POYO-MP model, and to single-session CEBRA, for which we ran a separate parameter sweep on the output dimensionality. We did not sweep parameters independently for each session, but we tuned the hyperparameters on a representative subset of 10 sessions and applied the best-performing configuration across all 99 sessions.

##### Scaling experiment

We computed velocity R^2^ for models trained on 1, 2, 5, 10, 20, 40, 60, 80 and 99 sessions, with 5 runs per training-set size and random session resampling for each run, except for the full 99-session set. Hyperparameters used were the same one as for the full dataset model (Supplemental Table S11).

##### Training cost

We compare the training cost of Unified CEBRA with CEBRA (21) and POYO-MP (39) (Supplemental Table S4).

- *Training Time*: The wall-clock time from the start of training to the saving of the model. For single-session CEBRA, it is the sum over the 99 single-session models. All models we trained ran on a single NVIDIA RTX A5000 GPU (24 GB). For POYO-MP, we report the training time given by Azabou et al. (39), obtained on 8 NVIDIA A40 GPUs (48 GB each). Because the hardware differs, we also report GPU-hours (number of GPUs *×* training time).
- *Parameters*. The number of trainable parameters, in millions (M). For POYO-MP, it was inferred from the scaling analysis and model sizes reported by Azabou et al. (39).
- *Forward FLOPs*. The number of floating-point operations for one forward pass on one sample, in gigaFLOPs (G). A sample is the smallest input that yields one embedding timestep: batch size 1 and an input window equal to the receptive field of the encoder. We counted the FLOPs of all convolutional and linear layers, and of the attention products for transformer models, counting each multiply-accumulate as 2 FLOPs. For CEBRA, we report the mean over sessions.
- *Training compute*. The total number of floating-point operations over training, in petaFLOPs (PFLOPs). A backward pass costs about twice a forward pass, so one training sample costs three times the forward FLOPs. Each CEBRA training step processes three samples per batch element (reference, positive and negative), giving a training compute of 3 *×*FLOPs*×* 3 *×B ×S*, with *B* the batch size and *S* the number of training steps. For CEBRA, it is summed over encoders. Unlike training time, training compute does not depend on the hardware.

#### Mouse sensorimotor Dataset

Mice learned a joystick task and performed a reach to grasp to center-in reaching movement. Layer 2/3 neurons across somatosensory and motor forelimb regions were recorded with 2P-imaging across 22 sessions from 6 mice. We used the baseline sessions presented in DeWolf et al. (43) using a nested, 3-fold cross-validation split.

We adopted the selection of regressors by DeWolf et al. (43), specifically the 3D hand position and velocity, as well as the 3D wrist position, for training the embedding. We fit five Unified CEBRA embeddings per dataset split and used auxiliary variables for the 3D hand and wrist position extracted with DeepLabCut (76, 77). DeWolf et al. (43) also provide a selection scheme for regressors from six different coordinate frameworks which were computed from the same underlying movement using a biome-chanical model of the mouse forearm. Besides 3D position and velocity (group 1), groups 2–6 cover biomechanical variables derived from DeWolf et al. (43): joint angles (group 2), joint torques (group 3), muscle force (group 4), muscle length/velocity (group 5), and spindle/Golgi afferent fiber activity (group 6). From the estimated embeddings, we fit linear regression models and estimate the R^2^ score of regressing from levels encoding lower level features (i.e., proprioceptive information in group 6 and muscle length/velocity in 5) to levels encoding higher level features (i.e., 3D position/velocity or angular position/velocity of the joints). For this decoding analysis, the test set embedding is computed by first aligning neural data based on the labels of the “source group”. The linear regression model is fitted on the train embedding, and evaluated on the test embedding using labels from the “target group”.

#### Hippocampus Dataset

This dataset (12) was previously used for benchmarking (20, 21) and contains 4 sessions of rats navigating on a linear track. In brief, bilaterally implanted silicon probes recorded multicellular electrophysiological data from CA1 hippocampus areas each from four male Long-Evans rats, while running on a 1.6-m-long linear track where they were rewarded with water at each end of the track. Sessions contain respectively 120, 48, 55, and 66 neurons, at Δ*t* = 25 ms. We kept the last 30% of the data for evaluation on each session; to note, this is different than how train/test was presented in Schneider et al. (21), thus the numbers are not directly comparable. Data pre-processing was as in Zhou and Wei (20). More dataset details can be found in Grosmark and Buzsáki (12).

For each trained model, we measured the cross-session consistency, providing insights into the similarity of latent spaces and the consistency of the model. We investigated the learned latent space behavioral decoding capabilities for position and direction on held-out data (20, 21). We trained a kNN decoder (k=36) on the training set outputs and tested on the held-out data. We also evaluated the held-out goodness-of-fit of the encoders, higher values indicating better data fitting.

#### Partial behavioral coverage

When recording from different animals or sessions, the behavior is not identical. Thus, it could be that some animals/sessions only cover a specific aspect of the behavioral space. To show that our method handles sessions with varying behavioral overlaps, we artificially cropped the training datasets for all four animals. Each session covered 25, 37.5, 50, 62.5, or 75% of the behavior space for an average behavioral overlap between sessions of respectively 0, 22, 42, 63, and 78%. For each run, the behavioral intervals defined in Supplemental Table S8 were randomly partitioned across the four sessions to assemble the training set, while the test set remained uncropped. Following model training, session-indexed output embeddings were concatenated into a unified global embedding. For CEBRA, we aligned the embeddings using Procrustes alignment (59). Finally, we trained a kNN decoder (k = 36) on the global embeddings to predict the position and direction on the uncropped held-out data (original data).

#### Place cells position attribution

We trained five regularized Unified CEBRA models using the same contiguous train-test split (70-30%). All hyperparameters were similar to the masked models, and the regularization weight was uniformly sampled from [0.05, 0.15] around a grid search optimum of 0.1 (empirical mean 0.101 *±*0.018). To test whether our feature attribution map **M** recovers known single-neuron coding, we computed position attributions for neurons across all four rats, whose place fields provide a ground truth. For each of the five unified models and each rat, a ridge readout **W** was fitted on the training split from the embedding to z-scored position and running direction (*α* = 1). The attribution **M**(*t*) = **J**(*t*)^*⊤*^**W** was computed on the validation split (last 30%), with **J**(*t*) averaged over the encoder’s input window, and restricted to the rat’s own neurons (other sessions were aligned to get the session-indexed embedding but each neuron was only considered once, for its session).

The place field peak was taken as the maximum of the neuron’s occupancy-normalized rate map, computed on the training split. First, position was binned to 0.02 m, and running directions were pooled. For a place cell, increasing its activity should move the decoded position toward its place field: forward when the animal is before the field, backward when it is past it. Its position attribution along the track should therefore change sign, from positive to negative, at the field. For each neuron, we averaged the position attribution also within 0.02 m bins for each model, then across models, pooling both running directions. We then located the main positive-to-negative zero crossing of this profile without using the rate map. To do this, each such crossing (excluding those within 0.08 m of track end) was scored by the summed magnitude of the contiguous positive (below) and negative (above) regions on either side, and the highest-scoring one was kept (linear interpolation within the bin). Neurons with no firing were excluded, and neurons with a rate peak within 0.25 m of a track end (approximate mouse size) were excluded from the statistics. 283 of 289 neurons have activity (training split); 179 have their peak 0.25–1.35 m from the track ends and enter the statistics; 178 of those have a crossing. We compared crossings with rate peaks by Pearson correlation over cells with a computed crossing. The null distribution was obtained by circularly shifting each rat’s attribution time series relative to behavior (2,000 shifts), recomputing the profiles and crossings, and comparing them with the true rate peaks and we reported the empirical one-sided *p*-value.

#### Two-photon Visual Coding Dataset

In the two-photon (2P) visual coding dataset of the Allen Brain Observatory, mice passively viewed movies. We focused on ten repeats of a monochrome movie (“Natural Movie 1”; 900 frames per repeat). Excitatory and inhibitory neurons in three visual cortical areas – the primary visual cortex (VISp) and two higher visual areas, VISl and VISpm – were recorded with 2P calcium imaging at Δ*t* = 33 ms, aligned to the movie frames, which were shown at 30 Hz (13). Inhibitory neurons were identified through three Cre-driver lines, Vip, Sst, and PV. Interneurons are sparse in each recording, so they were pooled across all available sessions of each Cre line. We selected sessions via the per-session Cre-line summary provided by the Allen Brain Observatory, and data processing is as in (78). To give excitatory and inhibitory sessions comparable sizes, each excitatory session was subsampled at random to match the distribution of inhibitory session sizes (exact number of neurons, sessions and mice used in Supplemental Table S13). More details on the dataset can be found at https://portal.brain-map.org/circuits-behavior/visual-coding-2p.

We trained a regularized Unified CEBRA encoder shared across all sessions, through a repeat-wise three-fold cross-validation: in each fold, one movie repeat (repeat 3, 6 or 9) was held out for testing, and three encoders with different random seeds were trained on the remaining repeats, for nine models in total. The features of the DINO model (79), as computed in (21), were used as auxiliary variables to guide training of the encoders; these features were extracted using the pretrained ViT/8 vision transformer DINO model that embeds video frames into a 768-dimensional feature space. Each session comprised ten repeats of the 900-frame movie. Regularization weight was uniformly sampled from [0.02, 0.08] around a grid-search optimum of 0.06 (0.0655*±* 0.0026 across runs). We reported the training cross-runs consistency and training goodness-of-fit (R^2^ = 0.985*±* 0.002, GoF: 9.199*±* 0.001; Supplemental Figure S3a-b). We evaluated each learned latent space by decoding frame identity on a held-out repeat. A kNN decoder (k = 36) was trained on the embedding of the nine training repeats and tested on the embedding of the held-out repeat, and we report accuracy within a 1-second window (97.3%). We also decoded six features extracted from the movie (see below) from the same embeddings with kNN decoders, none of which was used as the auxiliary variable (mean R^2^ = 0.88; Supplemental Figure S3c). Attribution maps, neuron rankings and decoders were computed from the training repeats of each fold only, and all results were evaluated on the corresponding held-out repeat and averaged over folds.

##### Excitatory-only Unified CEBRA control

To test whether inhibitory neurons contribute to the shared latent space, we trained the same unified model on excitatory neurons only. The number of neurons was matched exactly to the main model and, as only 220 excitatory sessions exist, the control used all of them. Session sizes followed the main model’s distribution within each area and the neurons kept in each were drawn at random. All other settings were identical to the main model. The control models were trained in the same three cross-validation folds, with two runs per fold (six models total).

For each feature, held-out decoding of the excitatory-only unified models was significantly lower than decoding with main unified models (permutation test stratified by cross-validation fold; 1,000 permutations for the excitatory-only control; Benjamini-Hochberg correction; *p <* 0.01; Supplemental Figure S3c).

##### Pseudo-session single-session CEBRA control

Because the movie is identical and frame-aligned across sessions, all sessions can also be combined without the unified sampler. Similar to (21), we stacked the same 2,924 neurons from the 234 sessions along the neuron axis at matching movie frames, forming a deterministically-aligned pseudo-session. A single-session regularized model with the same cross-validation, architecture, regularization and hyperparameters as the unified model was trained on this pseudo-session population, so that no cross-session sampling takes place, and we compared the same evaluation metrics.

The pseudo-session models produced similar decoding performances (Supplemental Figure S3c). The attribution map obtained were similarly reproducible across independently trained runs at the neuron and class levels (Supplemental Table S14), while ablating neurons based on its ranking affected decoding for the specific feature as much as those ranked by the unified map up to 2% of neurons and less at larger set sizes (Supplemental Figure S3e). Overall, those results indicated that the unified sampling does not distort class-level attributions.

##### Extra movie features

We decoded six low-level visual features from the latent embeddings: motion energy (Farneback optical flow), vertical and horizontal edge energies (Gabor filters, *f* = 0.1), luminance (mean frame intensity), and low-vs. high-spatial-frequency energy (2D Fourier spectrum split at radius *r* = 15).

## Supplemental Figures

**Figure S1.**
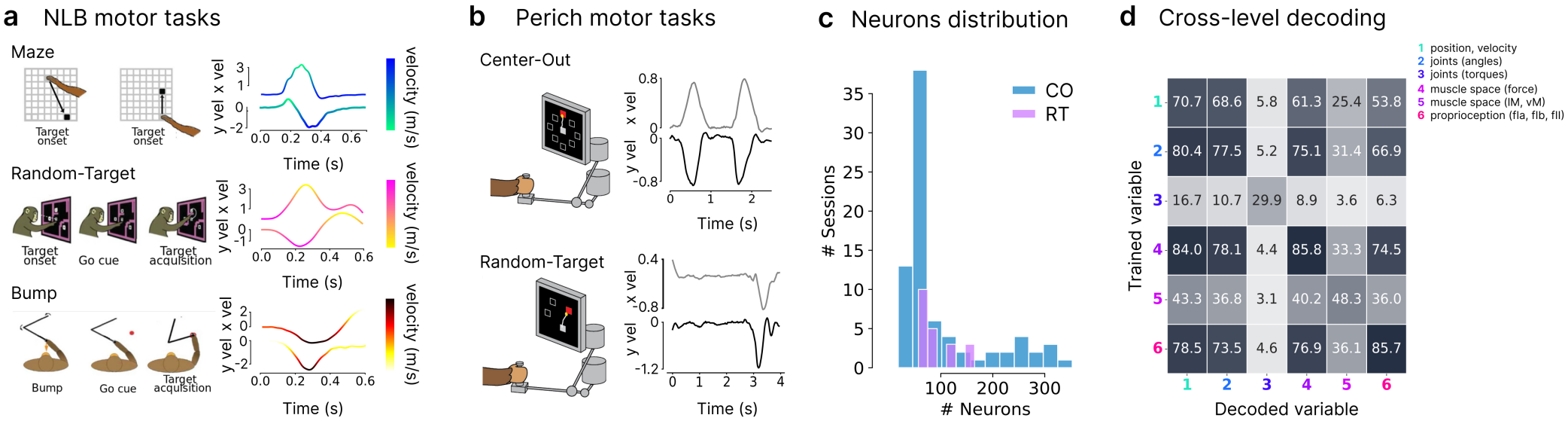
Framework, NLB and Perich datasets extended results. **a:** NLB tasks, taken from (10), and corresponding behavioral traces (x and y velocity) for a trial example. **b:** Perich tasks, Center-out (CO; 78 sessions) and random-target (RT; 21 sessions) reaching tasks (99 sessions, three monkeys; schematics adapted from (39)), with representative single-trial velocity profiles (right). **c:** Neurons per session (CO: 100.27 *±*9.51, 7,821 in total; RT: 90.67*±* 7.88, 1,904 in total; 9,725 neurons). **d:** Cross-level decoding R^2^ (%) confusion matrix. A model is trained on one representation (rows, Trained variable) and evaluated on another (columns, Decoded variable). Variables correspond to: (1) position and velocity, (2) joint angles, (3) joint torques, (4) muscle force, (5) muscle length/velocity, and (6) proprioception (fIa, fIb, fII). Darker cells indicate higher decoding performances.

**Figure S2.**
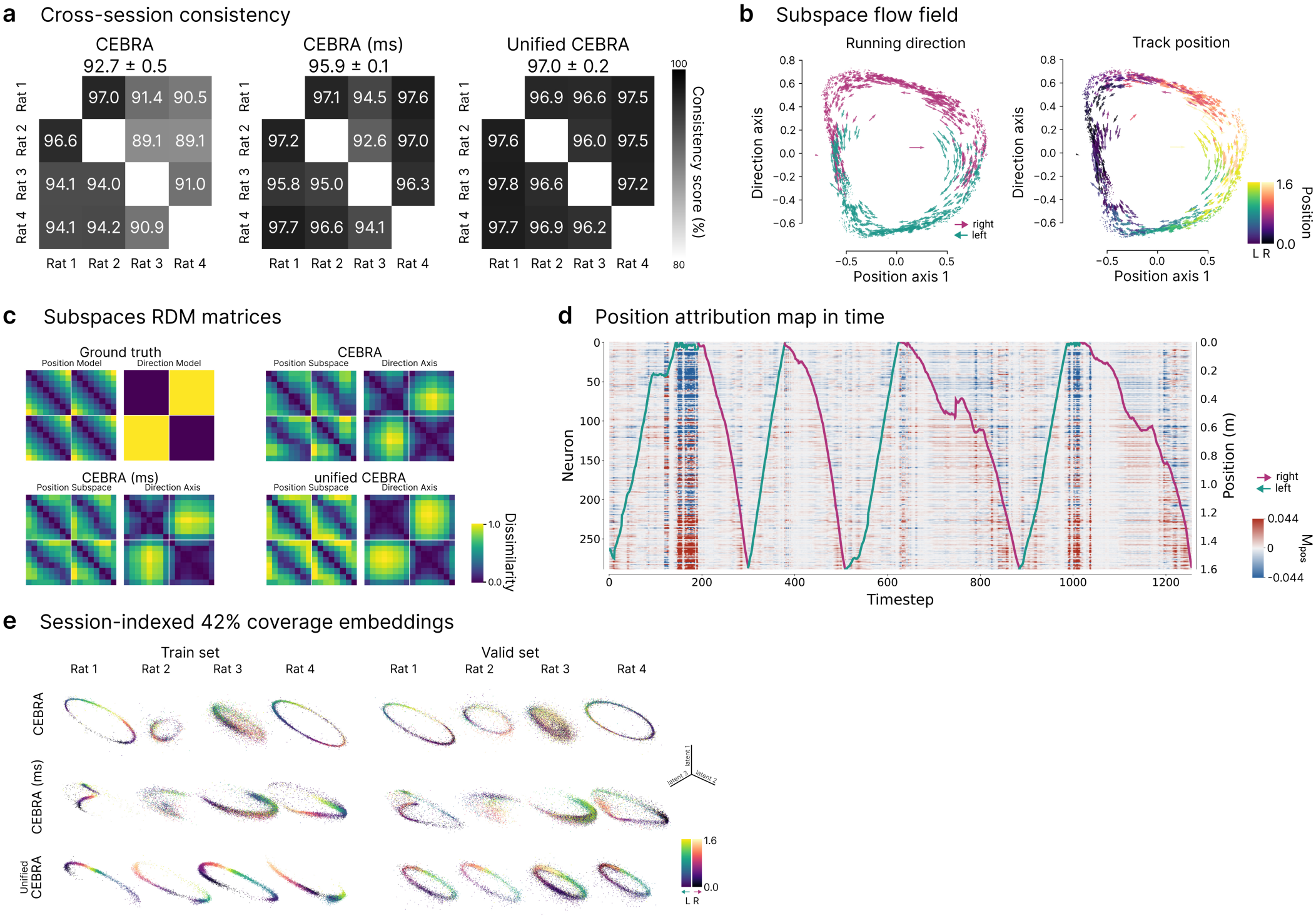
Hippocampus dataset extended results. **a:** Cross-session consistency scores. Embeddings from the single session encoders are aligned using Procrustes alignment (59) before computing the consistency. Values above the matrices: mean*±* SEM; n = 10 runs. **b:** Representational dissimilarity matrices (RDMs) across conditions (8 position bins × 2 running directions). Ground truth: RDMs predicted by position and direction. Per model: RDMs computed from the position subspace and direction axis of one example run, four rats pooled, on held-out data. CEBRA baselines aligned with Procrustes alignment (59) **c:** Direction axis versus first position axis (via subspace decomposition; Methods), with flow-fields highlighting dynamics. Left: colored by running direction, right: colored by position. **d:** Position attribution (**M**_Pos_) of each neuron over time across all sessions (validation split example; neurons sorted by place field peak). Red: increasing the neuron’s activity moves the decoded position forward along the track; blue: backward. Overlaid: position colored by direction. **e:** Session-indexed training and validation sets embeddings for 42% average overlap across sessions at training (3 out of 32 dimensions; colored by position and direction).

**Figure S3.**
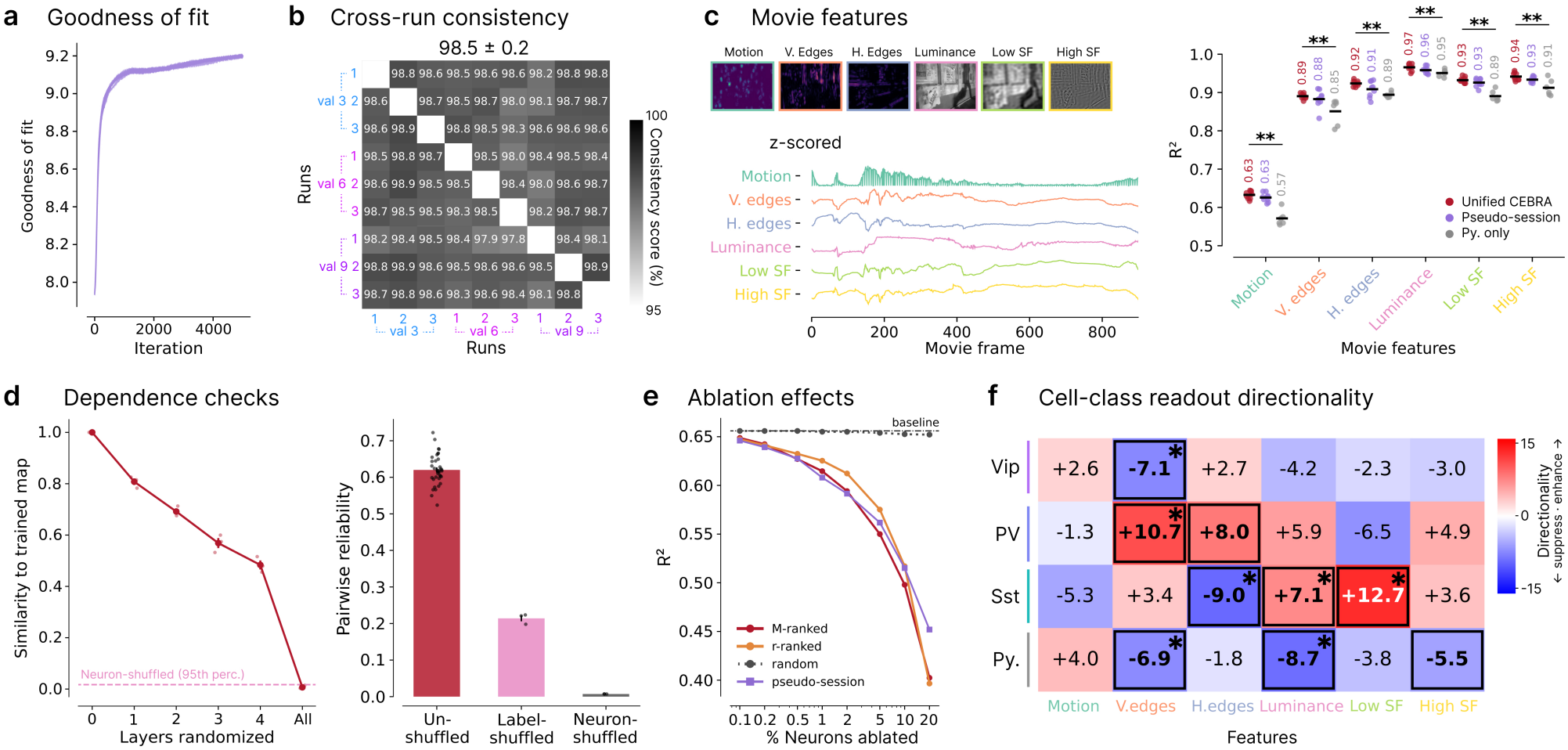
2P Visual Coding (Allen Institute) dataset extended results. **a:** Training goodness-of-fit (in bits) for the 9 model runs. **b:** Cross-run consistency scores matrix across the different folds (held-out on repeat 3, 6, or 9), and runs per fold (3 each). **c:** Movie features and decoding from the global latent embedding. Left, z-scored feature values across movie frames. Right, kNN decoding performance (k = 36; R^2^) per movie feature from the unified and pseudo-session latent embeddings (9 runs each) compared to a Unified CEBRA setup trained only on excitatory neurons (2,924 neurons, all 220 available sessions; 6 runs); individual dots: individual sessions averaged across runs; black bars and annotations: mean across sessions. Permutation test within cross-validation folds, Py + interneurons vs. Py-only Unified CEBRA, per feature, BH correction, *p <* 0.01. **d:** Dependence checks. Left, map similarity mean *±*SEM (3 runs) to the fully trained model as encoder layers are randomized cumulatively from the output layer inward; pink: neuron-shuffled null (95th percentile). Right, Pairwise map reliability for unshuffled models (9 runs, 36 pairs), label-shuffled controls (3 runs, 3 pairs), and a neuron-shuffled null. Black dots: individual comparisons. **e:** Decoding of each feature (R^2^) as a function of the percentage of neurons removed, ranked by attribution magnitude 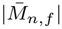 (M-ranked, red), marginal tuning |*r*_*n,f*|_ (r-ranked, orange), the pseudo-session model’s map (purple), or at random (gray, dotted). Dashed line: decoding with all neurons (baseline). Mean over the six features. **f:** Cell-type readout directionality. Percentage of readout-enhancing (positive-signed attribution) neurons relative to the unweighted mean across classes (blue = negative readout, red = positive readout). Boxed : Benjamini-Hochberg FDR *p <* 0.05 (session mixed model, parametric bootstrap, class-vs-mean contrasts; 24 tests). Stared: Agreement with marginal tuning directionality.

## Supplemental Tables

**Table S1.** Held-out Goodness-of-Fit (GoF; in bits). Values are comparable only within each dataset only, as goodness-of-fit depends on batch size and dataset. Mean *±* SEM over runs, with rats or sessions averaged within each run (hippocampus, NLB: 10 runs; Perich: 5 runs).

| **Dataset** | **CEBRA** | **CEBRA (ms)** | **Unified CEBRA** |
|---|---|---|---|
| NLB | 1.051 ± 0.002 | 0.898 ± 0.003 | 1.756 ± 0.006 |
| Perich | 0.563 ± 0.001 | — | 4.487 ± 0.007 |
| Hippocampus | 0.483 ± 0.003 | 0.463 ± 0.003 | 1.102 ± 0.003 |

**Table S2.**
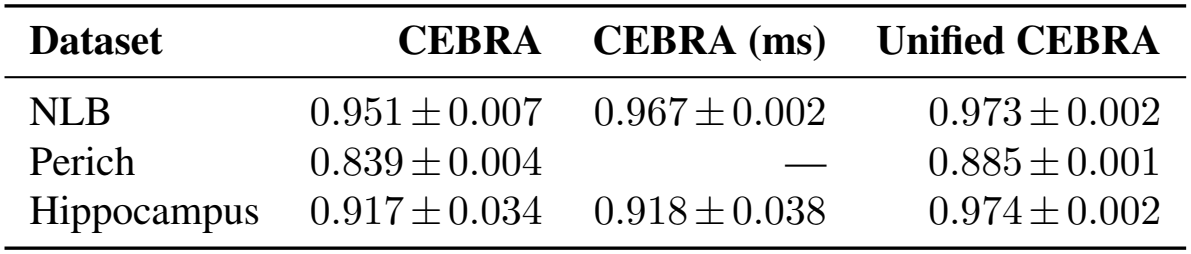
Training cross-run consistency (R^2^). Mean*±* SEM over runs, with rats or sessions averaged within each run (hippocampus, NLB: 10 runs; Perich: 5 runs).

**Table S3.** NLB temporal neuron and auxiliary labels shuffling control experiment. Velocity decoding performance (R^2^) pooled across datasets under temporal neuron or auxiliary variable shuffling at inference (encoder trained on intact data; evaluated on shuffled data) or training (encoder trained on shuffled data; evaluated on intact data). Auxiliary labels and label-guided matching left intact across conditions. *All but one:* each dataset kept intact in turn, with the other five shuffled; mean over datasets. Values report mean *±*SEM across runs (n = 10 for intact baselines and inference shuffle; n = 5 for training shuffles).

| **Condition** | **Shuffled modality** | **Shuffled sessions** | **Velocity R<sup>2</sup> (↑)** |
|---|---|---|---|
| CEBRA | — | — | 0.669 ± 0.002 |
| CEBRA (ms) | — | — | 0.642 ± 0.001 |
| Unified CEBRA | — | — | 0.926 ± 0.001 |
| Shuffled at *inference* | Neural activity | All | −0.062 ± 0.009 |
|  |  | All *but one* | 0.101 ± 0.004 |
|  | Auxiliary label | All | 0.113 ± 0.006 |
|  |  | All *but one* | 0.107 ± 0.005 |
| Shuffled at *training* | Neural activity | All | −0.054 ± 0.036 |
|  | Auxiliary label | All | 0.108 ± 0.022 |

**Table S4.** Model complexity, training compute and training time on the Perich dataset (99 sessions). Parameters: trainable parameters (millions, M). Forward FLOPs: floating-point operations for one forward pass on one sample (gigaFLOPs, G). Training compute: total floating-point operations over training (petaFLOPs, PFLOPs). Training time: wall-clock time on the listed hardware; GPU-hours: number of GPUs*×* training time. For CEBRA, parameters, training compute, training time and GPU-hours are summed over the 99 single-session models, and forward FLOPs are averaged over them. *\**: reported or inferred from Azabou et al. (39).

| Model | Parameters (M) | Forward FLOPs (G) | Training Compute (PFLOPs) | Hardware | Training Time | GPU-Hours |
|---|---|---|---|---|---|---|
| **Unified CEBRA** | 3.39 | 0.230 | 0.529 | 1× RTX A5000 | 17’06” | 0.3 |
| POYO-MP [39] | ∼12<sup>*</sup> | — | — | 8× A40<sup>*</sup> | ∼2 days<sup>*</sup> | ∼384 |
| CEBRA [21] | 90.93 | 0.037 | 34.090 | 1× RTX A5000 | 2h18’ | 2.3 |

**Table S5.** Velocity decoding R^2^ on Neural Latent Benchmark (NLB) datasets. For CEBRA baselines, POYO and Unified CEBRA, values report mean *±*SEM (n=10 runs). *EvalAI leaderboard:* Results taken directly from the official EvalAI leaderboard evaluated on private test split to contextualize current state-of-the-art performance. *Public evaluation split:* evaluated on publicly available held-out validation splits. *\**: trained on all six datasets.

|  | **Dataset & Velocity R<sup>2</sup> (↑)** |  |  |  |  |  |
|---|---|---|---|---|---|---|
| **Method** | MC-Maze | MC-Maze-S | MC-Maze-M | MC-Maze-L | MC-RTT | Area2-Bump |
| *EvalAI leaderboard* |  |  |  |  |  |  |
| Ctrl-TNDM [80] | 0.9362 | — | — | — | 0.6705 | 0.8950 |
| MINT [5] | 0.9121 | 0.8711 | 0.8958 | 0.9330 | 0.6559 | 0.8877 |
| AESMTE3 [73] | 0.9114 | 0.8669 | 0.9019 | 0.9252 | 0.6334 | 0.8999 |
| NDT-U [73] | — | 0.8490 | 0.9026 | 0.9157 | — | 0.8701 |
| NDT-L [73] | 0.8938 | — | — | — | 0.5895 | — |
| NeuralRoBERTa+RNNF [81] | — | 0.8254 | 0.8762 | 0.9185 | 0.6489 | 0.8761 |
| S5 [82] | 0.9043 | — | — | — | 0.6450 | 0.8727 |
| AutoLFADS [72] | 0.9097 | 0.7982 | 0.8680 | 0.9178 | 0.6167 | 0.8492 |
| STNDT [83] | 0.9095 | — | — | — | 0.6352 | 0.8937 |
| *Public validation split* |  |  |  |  |  |  |
| Neural kNN | 0.5732 | 0.3781 | 0.4985 | 0.4996 | 0.3549 | 0.7728 |
| POYO<sup>*</sup> [39] | 0.9307 ± 0.0006 | 0.8513 ± 0.0052 | 0.8857 ± 0.0031 | 0.9163 ± 0.0010 | 0.5085 ± 0.0053 | 0.8521 ± 0.0028 |
| CEBRA. [21] | 0.8550 ± 0.0012 | 0.4891 ± 0.0078 | 0.6541 ± 0.0036 | 0.7718 ± 0.0023 | 0.4756 ± 0.0024 | 0.7696 ± 0.0018 |
| CEBRA (ms)<sup>*</sup> [21] | 0.8002 ± 0.0009 | 0.4127 ± 0.0073 | 0.6300 ± 0.0034 | 0.7469 ± 0.0019 | 0.4916 ± 0.0016 | 0.7729 ± 0.0024 |
| **Unified CEBRA**<sup>\*</sup> | 0.9398 ± 0.0007 | 0.9439 ± 0.0009 | 0.9364 ± 0.0007 | 0.9462 ± 0.0007 | 0.8573 ± 0.0023 | 0.9332 ± 0.0014 |

**Table S6.**
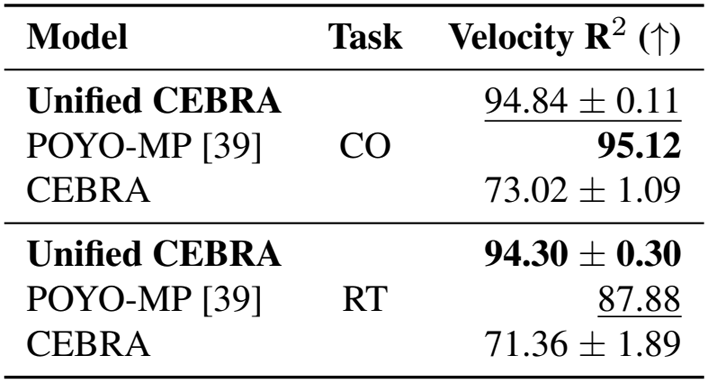
Velocity decoding performance on the Perich dataset. Mean velocity R^2^ across all sessions for the center-out (CO) and continuous random target (RT) acquisition tasks (n = 5 runs). Results for POYO are from Azabou et al. (39). multi-session CEBRA was limited by memory at that scale.

| **Model** | **Task** | **Velocity R<sup>2</sup> (↑)** |
|---|---|---|
| **Unified CEBRA** | CO | 94.84 ± 0.11 |
| POYO-MP [39] |  | **95.12** |
| CEBRA |  | 73.02 ± 1.09 |
| **Unified CEBRA** | RT | **94.30 ± 0.30** |
| POYO-MP [39] |  | 87.88 |
| CEBRA |  | 71.36 ± 1.89 |

**Table S7.** Separation of behavioral variables in the latent space. Double dissociation (DD) between the subspaces encoding two behavioral variables: reach direction and speed (both derived from hand velocity, the auxiliary variable) for the primate reaching datasets, and position and running direction for the hippocampus dataset. DD equals 2 when each subspace reflects only its own variable. Mean*±* SEM over runs, with rats or sessions averaged within each run (hippocampus, NLB: 10 runs; Perich: 5 runs). In every run, DD exceeded the 95th percentile of a label-shift null. —: not available (multi-session training failed on this dataset).

| **Dataset** | **Labels** | **CEBRA** | **CEBRA (ms)** | **Unified CEBRA** |
|---|---|---|---|---|
| NLB | Reach direction, Speed | 1.23 ± 0.01 | 1.22 ± 0.01 | **1.59** ± **0.01** |
| Perich | Reach direction, Speed | 0.41 ± 0.01 | — | **1.25** ± **0.04** |
| Hippocampus | Position, Direction | 1.61 ± 0.01 | 1.72 ± 0.01 | **1.94** ± **0.01** |

**Table S8.**
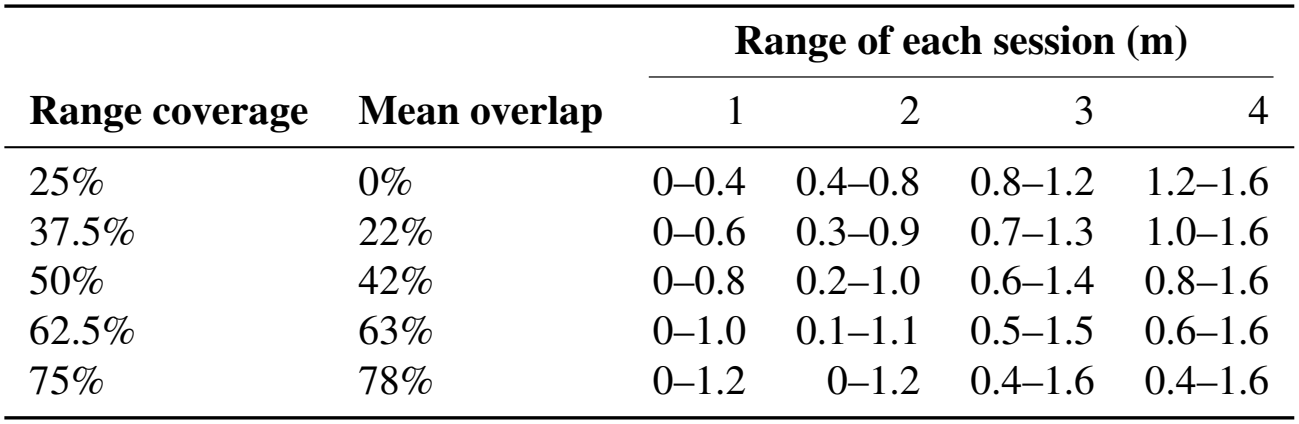
Behavioral ranges in the partial-coverage experiment. Each of the four rats was assigned one range of the 1.6-m track, assignments randomized across runs. *Range coverage:* track fraction covered by each session. *Mean overlap:* track length shared by two sessions’ ranges, averaged over the six pairs, expressed as a fraction of the range length.

| **Range coverage** | **Mean overlap** | **Range of each session (m)** |  |  |  |
|---|---|---|---|---|---|
|  |  | 1 | 2 | 3 | 4 |
| 25% | 0% | 0–0.4 | 0.4–0.8 | 0.8–1.2 | 1.2–1.6 |
| 37.5% | 22% | 0–0.6 | 0.3–0.9 | 0.7–1.3 | 1.0–1.6 |
| 50% | 42% | 0–0.8 | 0.2–1.0 | 0.6–1.4 | 0.8–1.6 |
| 62.5% | 63% | 0–1.0 | 0.1–1.1 | 0.5–1.5 | 0.6–1.6 |
| 75% | 78% | 0–1.2 | 0–1.2 | 0.4–1.6 | 0.4–1.6 |

**Table S9.** Hippocampus temporal neurons shuffling control experiment. Behavioral decoding performance on hippocampal recordings pooled across all four rats when neural activity is shuffled in time, per neuron, either at inference (encoder trained on intact data; evaluated on shuffled data) or at training (encoder trained on shuffled data; evaluated on intact data). Auxiliary labels and label-guided matching are left intact across conditions. *All but one:* each rat kept intact in turn, with the other three shuffled; mean over rats. Values report mean *±*SEM across runs (n = 10 for intact baseline and inference shuffle; n = 5 for training shuffles).

| **Condition** | **Shuffled sessions** | **Position R<sup>2</sup> (↑)** | **Direction R<sup>2</sup> (↑)** | **Position Error (↓)** |
|---|---|---|---|---|
| **CEBRA-Behavior** | — | 0.397 ± 0.004 | 0.293 ± 0.007 | 0.162 ± 0.002 |
| **Unified CEBRA** | — | 0.782 ± 0.005 | 0.748 ± 0.003 | 0.043 ± 0.000 |
| **Shuffled at inference** | All | −1.123 ± 0.032 | −0.924 ± 0.036 | 0.570 ± 0.009 |
|  | All but one | −0.553 ± 0.026 | −0.316 ± 0.025 | 0.381 ± 0.006 |
| **Shuffled at training** | All | −0.370 ± 0.086 | −0.088 ± 0.023 | 0.540 ± 0.017 |

**Table S10.** Hippocampus decoding performance across training paradigms and fine-tuning to held-out rats. CEBRA, CEBRA multi-session, and *De Novo* models are trained on all sessions. Fine-tuning models report adaptation performance on each held-out subject after pre-training on the remaining three rats. Values: mean *±*SEM across runs. Position error (Pos. Err.) is reported in meters (m); the mean column reports across-rat performance. Please note, this is a different data split than used in(21); see Methods.

| **Model** | **Phase** | **Metric** | **Rat 1** | **Rat 2** | **Rat 3** | **Rat 4** | **Mean** |
|---|---|---|---|---|---|---|---|
| CEBRA | — | Pos. R<sup>2</sup> (↑) | 0.749 ± 0.009 | 0.663 ± 0.007 | −0.067 ± 0.009 | 0.244 ± 0.012 | 0.397 ± 0.004 |
|  |  | Dir. R<sup>2</sup> (↑) | 0.435 ± 0.006 | 0.482 ± 0.012 | 0.137 ± 0.005 | 0.119 ± 0.015 | 0.293 ± 0.007 |
|  |  | Pos. Err. (↓) | 0.086 ± 0.001 | 0.120 ± 0.001 | 0.288 ± 0.004 | 0.155 ± 0.005 | 0.162 ± 0.002 |
| CEBRA (ms) | — | Pos. R<sup>2</sup> (↑) | 0.753 ± 0.010 | 0.693 ± 0.006 | −0.102 ± 0.011 | 0.175 ± 0.011 | 0.380 ± 0.004 |
|  |  | Dir. R<sup>2</sup> (↑) | 0.466 ± 0.009 | 0.472 ± 0.008 | 0.104 ± 0.006 | 0.161 ± 0.010 | 0.301 ± 0.004 |
|  |  | Pos. Err. (↓) | 0.076 ± 0.001 | 0.122 ± 0.001 | 0.267 ± 0.003 | 0.132 ± 0.002 | 0.149 ± 0.001 |
| **Unified CEBRA** | *De Novo*<br>(All 4 rats) | Pos. R<sup>2</sup> (↑) | 0.945 ± 0.003 | 0.958 ± 0.004 | 0.295 ± 0.018 | 0.930 ± 0.002 | 0.782 ± 0.005 |
|  |  | Dir. R<sup>2</sup> (↑) | 0.883 ± 0.002 | 0.928 ± 0.003 | 0.352 ± 0.008 | 0.879 ± 0.005 | 0.761 ± 0.004 |
|  |  | Pos. Err. (↓) | 0.039 ± 0.001 | 0.045 ± 0.001 | 0.043 ± 0.001 | 0.042 ± 0.000 | 0.043 ± 0.000 |
|  | Fine-tuning<br>(Held-out rat) | Pos. R<sup>2</sup> (↑) | 0.950 ± 0.006 | 0.960 ± 0.004 | 0.307 ± 0.009 | 0.934 ± 0.006 | 0.788 ± 0.000 |
|  |  | Dir. R<sup>2</sup> (↑) | 0.892 ± 0.011 | 0.901 ± 0.007 | 0.314 ± 0.012 | 0.887 ± 0.006 | 0.748 ± 0.003 |
|  |  | Pos. Err. (↓) | 0.043 ± 0.001 | 0.050 ± 0.001 | 0.046 ± 0.001 | 0.051 ± 0.002 | 0.047 ± 0.000 |

**Table S11.** Model parameters for CEBRA baselines and Unified CEBRA. When indicated, dropout (rate 0.1) was applied to all convolutional layers. Unified CEBRA parameters for Hippocampus and 2P Visual Coding determined via grid search for cross-run consistency, initialized from (21). For Perich, first value: Unified CEBRA, second: CEBRA; CEBRA multi-session was limited by memory at that scale. For 2P Visual Coding, Unified CEBRA and CEBRA were trained and evaluated, not CEBRA multi-session. *\**: NLB parameters applied for CEBRA multi-session and Unified CEBRA models; individual tuning for single-session CEBRA provided in Table S12.

|  | **Datasets** |  |  |  |  |
|---|---|---|---|---|---|
| **Parameters** | NLB* | Hippocampus | 2P Visual Coding | Perich | Sensorimotor |
| Iterations | 5000 | 5000 | 5000 | 2000 | 2500 |
| Receptive Field | 36 | 10 | 10 | 40 | 10 |
| Dropout | ✓ | ✗ | ✗ | ✗ | ✗ |
| Time Offsets | 4 | 10 | 10 | 3 | 10 |
| Batch Size | 256 | 2048 | 2048 | 128/512 | 1024 |
| Output Dimension | 45 | 32 | 128 | 32/16 | 6 |
| Hidden Units | 256 | 32 | 64 | 128 | 128 |
| Learning Rate | 8.39×10<sup>−5</sup> | 0.0003 | 10<sup>−5</sup> | 0.001 | 10<sup>−4</sup> |
| Temperature | 0.4241 | 1 | 1 | 0.2016 | 0.5 |

**Table S12.** Tuned hyperparameters for single-session CEBRA models across NLB sub-datasets. Only parameters deviating from the global baseline (Supplemental Table S11) are reported; identical configurations across Maze task scales were found (large, medium, small), hence grouped.

| **Parameters** | **Datasets** |  |  |  |
|---|---|---|---|---|
|  | Area2-Bump | MC-Maze | MC-Maze l/m/s | MC-RTT |
| Learning Rate | 8.39 × 10<sup>−5</sup> | 1.70 × 10<sup>−4</sup> | 4.66 × 10<sup>−5</sup> | 8.39 × 10<sup>−5</sup> |
| Temperature | 0.4241 | 0.0100 | 0.0381 | 0.4020 |

**Table S13.** Input data breakdown across visual areas and cell classes. Values are reported as neurons /sessions /mice. Distribution of two-photon calcium imaging recordings across primary (VISp) and higher visual areas (VISl, VISpm). One mouse can have recordings in multiple brain regions. Subpopulations comprise excitatory pyramidal neurons (Exc) and three Cre-targeted inhibitory interneuron subclasses (*PV, Sst, Vip*). *\**: VISpm lacks *PV* recordings due to dataset availability.

| **Visual Area** | **Excitatory** | ***PV*** | ***Sst*** | ***Vip*** | **Total** |
|---|---|---|---|---|---|
| VISp | 584 / 50 / 43 | 207 / 16 / 9 | 178 / 17 / 12 | 199 / 17 / 12 | 1,168 / 100 / 76 |
| VISl | 499 / 37 / 33 | 48 / 5 / 5 | 209 / 15 / 13 | 242 / 17 / 14 | 998 / 74 / 65 |
| VISpm | 379 / 30 / 27 | —<sup>*</sup> | 151 / 14 / 11 | 228 / 16 / 12 | 758 / 60 / 50 |
| **Total** | **1,462 / 117 / 87** | **255 / 21 / 9** | **538 / 46 / 29** | **669 / 50 / 24** | **2,924 / 234 / 149** |

**Table S14.** Pairwise cross-run reliability of attribution maps. Neuron level: mean *±*SEM Spearman rank correlation of |**M**_*n,f*|_ across all neurons, averaged over the features and runs. Class level: Pearson correlation of directional class biases across 24 class*×* feature entries. Attribution variants compose readout **W** with forward Jacobian **J** with a control derived from pseudo-session models.

| **Composition with W** | **Neuron** | **Cell-type** |
|---|---|---|
| **Forward J** | 0.620 ± 0.018 | 0.881 ± 0.017 |
| Pseudo-session map | 0.583 ± 0.009 | 0.931 ± 0.008 |

**Table S15.** Decoding after removing 5% of neurons (146 of 2,924). Held-out decoding performance (R^2^) of ridge regression decoders trained on raw activity: with all neurons (baseline), after removing the neurons ranked highest by the attribution map for each feature (M-ranked), and after removing size-matched control sets (mean over 10 draws). Parentheses: number of control draws with higher R^2^ than the M-ranked set, i.e., whose removal impaired decoding less. Mean row: mean R^2^ across the six features *±* SEM.

| **Feature** | **Baseline R<sup>2</sup>** | **M-ranked R<sup>2</sup>** | **Random-matched** | **Class × Area-matched** | **Firing-rate-matched** | **Depth-matched** | **Marginal tuning** |
|---|---|---|---|---|---|---|---|
| Motion | 0.458 | 0.378 | 0.458 (10/10) | 0.454 (10/10) | 0.457 (10/10) | 0.458 (10/10) | 0.383 |
| V. edges | 0.684 | 0.597 | 0.682 (10/10) | 0.666 (10/10) | 0.678 (10/10) | 0.680 (10/10) | 0.616 |
| H. edges | 0.647 | 0.557 | 0.642 (10/10) | 0.630 (10/10) | 0.642 (10/10) | 0.644 (10/10) | 0.577 |
| Luminance | 0.738 | 0.590 | 0.736 (10/10) | 0.727 (10/10) | 0.732 (10/10) | 0.736 (10/10) | 0.627 |
| Low SF | 0.661 | 0.527 | 0.660 (10/10) | 0.652 (10/10) | 0.661 (10/10) | 0.659 (10/10) | 0.563 |
| High SF | 0.749 | 0.651 | 0.746 (10/10) | 0.736 (10/10) | 0.746 (10/10) | 0.748 (10/10) | 0.686 |
| **Mean** | 0.656 ± 0.043 | **0.550 ± 0.038** | 0.654 ± 0.043 | 0.644 ± 0.042 | 0.653 ± 0.042 | 0.654 ± 0.043 | 0.575 ± 0.042 |

**Table S16.** Decoding across set sizes (0.1–20% of neurons; 3–585 neurons). Held-out decoding performance (R^2^, mean *±*SEM across the six features) with all neurons (baseline), after removing the neurons ranked highest by the attribution map (M-ranked), or control sets of the same size (mean over 10 draws): firing-rate-matched, random, and neurons ranked by marginal tuning (|*r*_*f*_ |; single deterministic set). Parentheses: number of features for which the M-ranked set had higher R than the control.

| **Set size** | **Baseline**<br>**R**<sup>2</sup> | **M-ranked**<br>**R**<sup>2</sup> | **Firing-rate-**<br>**matched** | **Random-**<br>**matched** | **Marginal**<br>**tuning** |
|---|---|---|---|---|---|
| 3 (0.1%) | 0.656 ± 0.043 | 0.649 ± 0.043 | 0.656 ± 0.043 (6/6) | 0.656 ± 0.043 (6/6) | 0.647 ± 0.042 (4/6) |
| 6 (0.2%) | 0.656 ± 0.043 | 0.642 ± 0.042 | 0.656 ± 0.043 (6/6) | 0.656 ± 0.043 (6/6) | 0.641 ± 0.043 (3/6) |
| 15 (0.5%) | 0.656 ± 0.043 | 0.627 ± 0.041 | 0.655 ± 0.043 (6/6) | 0.656 ± 0.043 (6/6) | 0.632 ± 0.043 (5/6) |
| 29 (1%) | 0.656 ± 0.043 | 0.615 ± 0.042 | 0.655 ± 0.043 (6/6) | 0.655 ± 0.043 (6/6) | 0.625 ± 0.044 (6/6) |
| 58 (2%) | 0.656 ± 0.043 | 0.594 ± 0.041 | 0.654 ± 0.043 (6/6) | 0.655 ± 0.043 (6/6) | 0.612 ± 0.045 (6/6) |
| 146 (5%) | 0.656 ± 0.043 | 0.550 ± 0.038 | 0.653 ± 0.042 (6/6) | 0.654 ± 0.043 (6/6) | 0.575 ± 0.042 (6/6) |
| 292 (10%) | 0.656 ± 0.043 | 0.498 ± 0.035 | 0.652 ± 0.043 (6/6) | 0.653 ± 0.043 (6/6) | 0.517 ± 0.043 (5/6) |
| 585 (20%) | 0.656 ± 0.043 | 0.402 ± 0.034 | 0.646 ± 0.041 (6/6) | 0.652 ± 0.042 (6/6) | 0.397 ± 0.040 (3/6) |

**Table S17.**
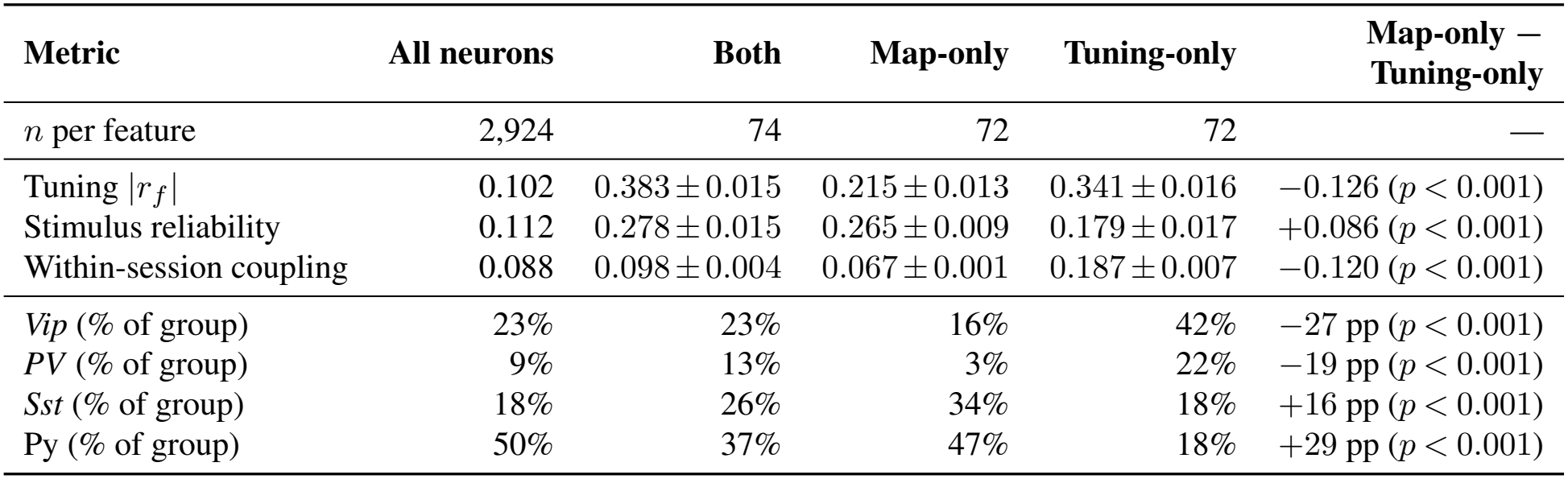
Properties of top-ranked neurons identified by attribution maps vs. marginal tuning. Comparisons across all neurons, intersecting units (both), and disjoint subsets (map-only vs. tuning-only) at 5% selection (146 neurons per feature; 74 overlapping, 72 disjoint). Values report mean *±*SEM across six features for continuous metrics, or percentage of group membership for cell classes. The difference column (map-only *−*tuning-only) reports raw differences and percentage points (pp) alongside two-sided *p*-values.

| **Metric** | **All neurons** | **Both** | **Map-only** | **Tuning-only** | **Map-only −<br>Tuning-only** |
|---|---|---|---|---|---|
| *n* per feature | 2,924 | 74 | 72 | 72 | — |
| Tuning \|*r*<sub>*f*</sub>\| | 0.102 | 0.383 ± 0.015 | 0.215 ± 0.013 | 0.341 ± 0.016 | −0.126 (*p* < 0.001) |
| Stimulus reliability | 0.112 | 0.278 ± 0.015 | 0.265 ± 0.009 | 0.179 ± 0.017 | +0.086 (*p* < 0.001) |
| Within-session coupling | 0.088 | 0.098 ± 0.004 | 0.067 ± 0.001 | 0.187 ± 0.007 | −0.120 (*p* < 0.001) |
| *Vip* (% of group) | 23% | 23% | 16% | 42% | −27 pp (*p* < 0.001) |
| *PV* (% of group) | 9% | 13% | 3% | 22% | −19 pp (*p* < 0.001) |
| *Sst* (% of group) | 18% | 26% | 34% | 18% | +16 pp (*p* < 0.001) |
| Py (% of group) | 50% | 37% | 47% | 18% | +29 pp (*p* < 0.001) |

## Notes

### Summary of Updates

the second authors name was misspelled on the website. the PDF is unchanged.

